# Closing the loop: tracking and perturbing behaviour of individuals in a group in real-time

**DOI:** 10.1101/071308

**Authors:** Malte J. Rasch, Aobo Shi, Zilong Ji

## Abstract

Quantitative description and selective perturbation of individual animals in a social group is prerequisite for understanding complex social behaviors. Tracking behavioral patterns of individuals in groups is an active research field, however, reliable software tools for long-term or real-time tracking are still scarce. We developed a new open-source platform, called **xyTracker**, for online tracking and recognition of individual animals in groups. Featuring a convenient Matlab-based interface and a fast multi-threading C++ core, we achieved an > 30× speed-up over a popular existing tracking method without loss in accuracy. Moreover, since memory usage is low, many hours of high-resolution video files can be tracked in reasonable time, making long-term observation of behavior possible. In a number of exemplary experiments on zebra fish, we show the feasibility of long-term observations and how to use the software to perform closed-loop experiments, where the tracked position of individuals is fed-back in real-time to a stimulus presentation screen installed below the fish-tank. Visual stimulation capabilities is incorporated into **xyTracker** and can be based on any behavioral features of all members of the group, such as, collective location, speed, or direction of movement, making interesting closed-loop experiments for investigating group behavior in a virtual reality setting possible.

## 1 Introduction

Classical behavioral research of animals has been traditionally limited by the lack of quantitative description of their complicated behavioral patterns, since raw observations had to be collected by “hand”. Modern advances of machine learning and computer vision allow to automate the description of behavior of many interacting animals on a much more quantitative computational level (e.g. [5, 57, 29]). Accurate and automated tracking of individual animals are at the core for quantitatively analyzing group behavior [21]. Automatic video tracking of individuals within a group of animals, such as a flock, shoal or school has only just begun and already yielded promising insights about the dynamics of groups of ducks [37], starlets [8], ants [40], mice[49], and fish [19, 26, 36, 15].

For the quantitative investigation of group behavior and learning, automatic tracking tools are still lacking. In particular, it would be desirable to perturb the behavior of some members of a group locally, e.g. using a visual stimulus which can be seen only by one individual of the group. Even better would be a visual stimulus presentation that could be “controlled” by the behavior of a subset of the individual in the group, to establish closed-loop feedback (learning) experiments for investigation of collective response patterns. This would extend both the associate learning paradigm done only for single individuals [50, 6, 42], and previously conducted single-individual closed-loop experiments, e.g. using a ight simulator of ies [14] or for investigating the ocularmotor reex of a single zebra fish larvae [10, 48]. Moreover, such a system would allow for controlled experimental verification of model predictions how swarms collectively response to local permutations [16].

Recently, considerably progress has been made in the area of multi-object tracking, an active discipline in computer vision. However, this research in computer vision is mainly focused on either tracking massive amounts of features with low importance on maintaining the individuality of objects (for a recent review see [17]), or on the problem of tracking humans in a cluttered environment (e.g. [59, 58, 33]). For behavioral research in ecology, only a selected number of available tracking systems of general purpose exists (for a recent review see [21]), and some solutions are specifically adopted for groups of fish [45, 25, 55, 41]. When tracking humans, individuals are often easily distinguishable and the challenge lies more in maintaining trajectories in a cluttered (3D) scene. On the other hand, the main difficulty of tracking groups of animals, such as fish, in a laboratory setting, is that individuals are often highly similar, so that close interaction or crossings of tracks results in intermingling of traces and erroneous mixing of individuals. Moreover, interactions happen frequently, are socially relevant [2, 24], and involve non-predictable, highly non-linear trajectories. Once two tracks are mixed, the error might not be corrected in later times and, and in the long-term, the identity of all individuals will be lost during tracking, that is, the identity of the individual tracked in the beginning will be diffierent from the identity tracked at the end of the observation period.

To solve this problem of the mixing of tracks, one approach is to extend the frameby-frame detection-based tracking of individuals by gathering information about their appearance. In [45], the authors propose to first track all detected objects (with a possible mixing of identity), then learn a classifier based on visual features of individuals in time periods where individuals are well distributed in space, and then, based on this classification, correct the wrong assignments of detection to tracks at times when individuals are very close nearby (crossing). While this method is successful in avoiding the accumulation of errors when tracking individual animals, a major drawback is that the method requires at least two passes through the recorded video file: one for tracking, the second for learning and classifier and classification of identities between crossing points. Thus, this method cannot be applied for online detection and identification in real-time.

We here present a new open-source software platform, **xyTracker**^1^, which is able to detect and follow individual animals in a group by online learning the identity of each animal from appearance features. Our system tracks in real-time and includes a stimulus presentation system for an affordable one-camera setup, which allows for closed-looped collective response experiments.

In our system, if individual tracks crossed recently, current identities of the tracks are compared to those before the crossing event and thus ensures that wrong assignments after crossings are corrected immediately by switching tracks in retrospect. Our system additionally implements a simple version of multiple hypothesis tracking [47, 30] to compute globally optimal trajectories in an online manner. Moreover, our tracking solution provides and easy way to present any kind of (visual) stimulus using the well-established PsychoToolbox [31] based on the current position of individual animals tracked in realtime with minimal lag (on the order of tens of milliseconds). While our software is mainly developed and tested for groups of fish, it is essentially parameter free^2^ as it learns appearance features and other parameters on the fly and is thus able to track any (cylindrical to oval-shaped) animals in a planar environment from above.

We implemented **xyTracker** in a performance optimized manner, by utilizing thread-based video grabbing and object detection (using the OpenCV library), but nevertheless providing an user-friendly Matlab interface. On a personal (quad-core) computer, videos having about 3 megapixel frames can be handled in less than 20 ms per frame, and a 7 minutes example video of behavior 5 fish is tracked in about 3:5 minutes. Moreover, since the Matlab interface is class-based, modules and functionality of xyTracker, such as the algorithm used for the classification or stimulus presentation, can be easily extended or exchanged by overloading existing classes.

In this paper we describe our software platform and compare the detection and tracking rate to available methods (in particular [45]). We show a number of experiments for validating the applicability of **xyTracker** for real-world behavioral studies and closed-looped experiments using groups of fish as an example.

## 2 Methods

The problem of tracking n individuals over time can be described by finding the spatial positions (e.g. pixel locations in a stationary video), **x***k*(*t*), where t is time (or frame number) and *k* = 1 … *n* the index of the *n* tracks. In our approach, this general problem can be roughly dissected into two steps per video frame: (1) Detection of all objects in the frame and (2) object-to-track assignment. For the first step, a frame is searched for any potential object *qi*. In the second step, a number of features such as the object pixel position, size and shape of each detection *qi* are extracted, and compared with previous features of existing tracks. Based on this comparison, detections are assigned to existing tracks.

Finally, when repeated for am frames, a trajectory is computed for each of the tracked animals. To illustrate the tracking results of **xyTracker**, trajectories of 5 fish are plotted together with extracted shape features in Fig. 1.

**Figure 1:**
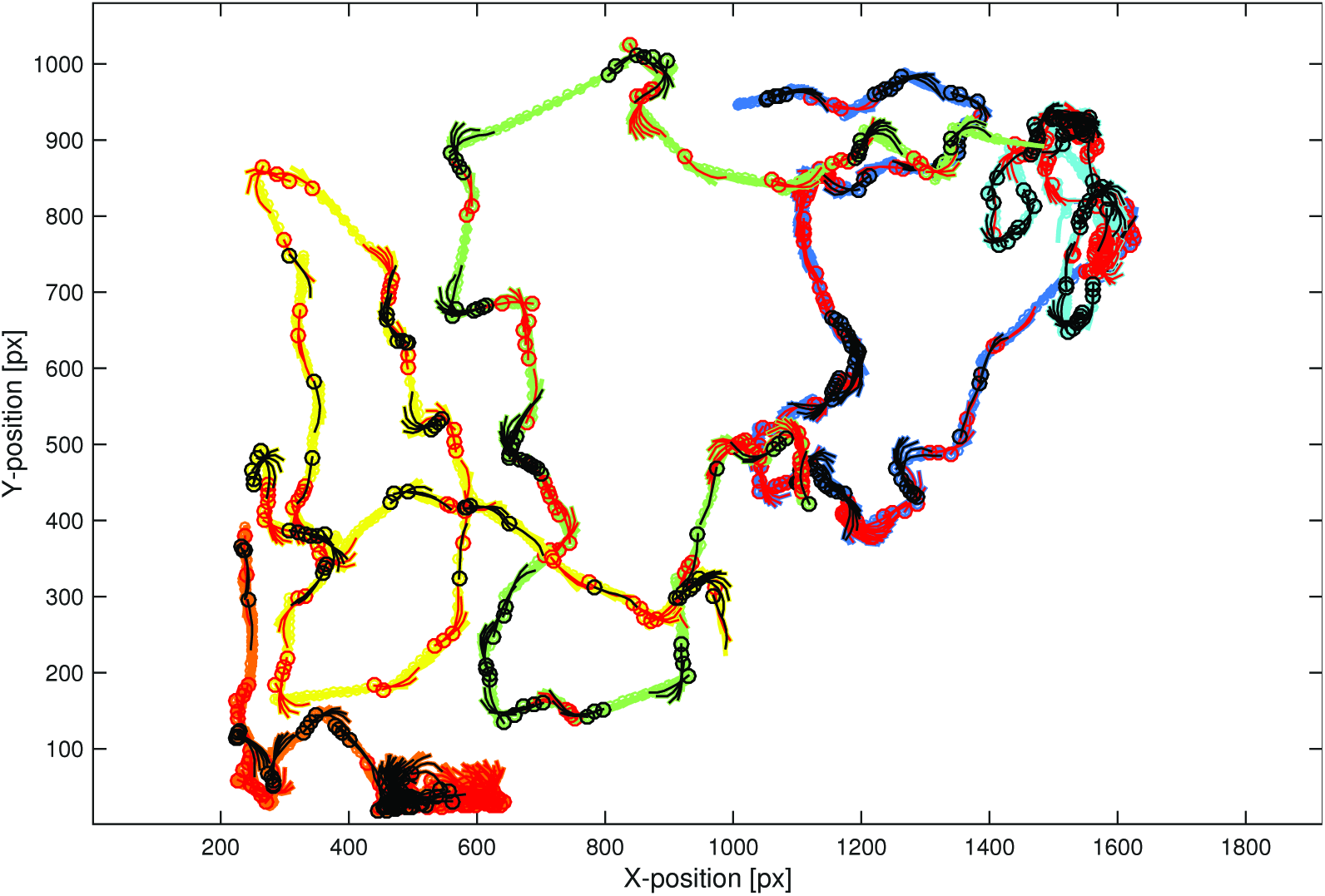
Example traces with shape features. Solid lines correspond to the mean position of all center line points per frame. If the center line bends (during turns) the whole center line is plotted in addition for each frame (red for rightward bends, black for leftward bends, a circle indicates the head position). One observes how fish bend their body to turn around. Also note that fish are either in a swimming or exploration state. The video is taken from [45].

In the following, we describe the object detection method, features and distances, and the detection-to-track assignments. Then, we describe the algorithms used for object classification and correction of potential wrong assignments, which is done in addition to the two steps above. For illustration of the tracking method in following, we use a video of 5 medaka fish provided with in the publication of [45]. However, our system can be applied to groups of any oval-shaped, lateral symmetric animal on a planar environment, with a camera mounted above (see Discussion). Although **xyTracker** works out-of-thebox for many problems, it is very flexible in setting parameters if necessary (see usage section). During the methods, we will therefore mention the name of the corresponding parameter options in the **xyTracker** implementation in footnotes.

### 2.1 Object detection

Let *I_t_*(*x*, *y*) be the gray scale intensity in the video frame at time *t* at pixel positions *x* and *y*. Colors, which can be included for the classification of identity of the tracked animals (see below), are not used during the object detection step^3^. For segmentation of moving objects from a static background, a common method is to threshold the intensity of the pixels in comparison to a background model and look for spatially connected patches of pixels above threshold. We found that the following simple method works very well in practice^4^,

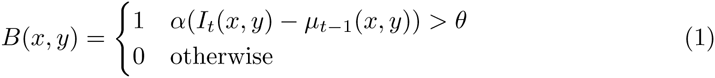

where *θ* is a threshold and *α* = -1 or *α* = 1 depending on whether light objects are in front of a dark background or vice versa^5^. The background average image *µ_t_*(*x*, *y*) is derived via leaky integration of the intensity image, *μ*_t_(*x*,*y*) = *μ*_*t*−1_(*x*,*y*)(1 − 1/*τ*_bckg_) + *I*_*t*−1_(*x*,*y*)/*τ*_bckg_, which is in practice computed only every *n*th frame to save computational time^6^. The time constant^7^ *τ*_bckg_ determines the time scale of variability of the background and is assumed to be much larger than the time scale of animals movements. We found that *τ*_bckg_ = 500 frames (about 20 seconds for our frame rate) is a reasonable setting which is robust to mild forms immobility in case of fish. However, this time constant depends on the application and might have to be adjusted^8^, e.g. when animals do note move for a long time because of freezing behaviors. The threshold *θ* is automatically determined during the first few tens of frames using the Otsu's method (in the OpenCV implementation) and then fixed for the remainder. The threshold can be adjusted within the **xyTracker** if detections are too small or too noisy^9^.

From the black-and-white mask image, *B*(*x*, *y*), “blobs” are extracted via a graph-theoretic approach for searching connected components, a common method in computer vision [22], again using the OpenCV toolbox or the Matlab equivalent. After applying some morphological closing and opening operations to remove noise pixels, each extracted blob is is principle regarded as a potential object, *q_i_*. The approximate body length, *L*, and approximate body width *W* of detected animals are used to calculate some bounds on the possible size of blobs. For example, if the area of a detection is smaller than a fraction or larger than a multiplicity of the expected area *LW*, a detection is discarded. Also the size of the identity features is based on these parameters (see below). They^10^ can be also given explicitly during startup of **xyTracker** if known, or are otherwise estimated automatically at the beginning.

#### 2.1.1 General blob feature extraction

From each detected blob *q_i_*, a number of feature are extracted. First, an ellipse is fitted to the shape of the blob, and the two half-axis lengths, orientation, and its center is used as features. An enclosing rectangular bounding box, and the number of the pixels in the blobs, and its area can also be used as features.

Moreover, using the bounding box, further features are extracted based on the pixel values belonging to the detected object. Since animals have often bilateral geometry, with mirror-axis parallel to the direction of movement (when viewed from above), we describe the shape of the blob with a simple “center line”, that is the mid-line of an cylinder-like body, that possibly bends (Fig. 2).

**Figure 2:**
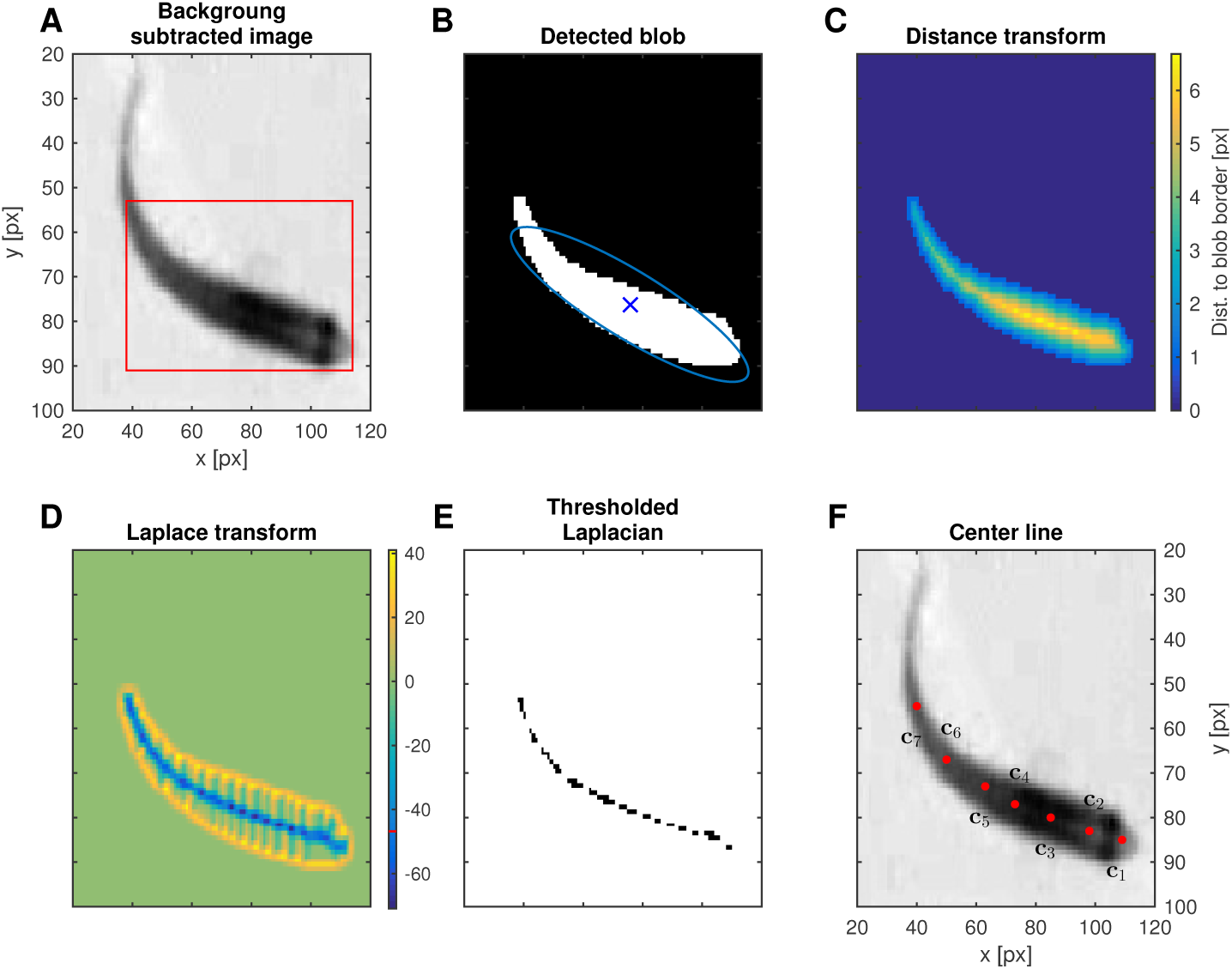
Extraction of features from detections. A: Raw image with subtracted background *I*(*x*, *y*) – *µ*(*x*, *y*). Bounding box of the detected blob (in B) plotted in red. B: Detected blob *q_i_* from the thresholded image *B*(*x*, *y*) after morphological transformation (dilation and erosion) that in this case removed noise and thin areas (like the tip of the tail) from the blobs. A number of feature *f_l_*(*q_i_*) are extracted from the detection *q_i_*, for instance an elliptic regions is fitted to the blob to extract its center, orientation, and axis lengths (blue line). C: Distance transform *d*(*x*, *y*) of the image in B. The value of the transform is equal to the distance of pixels to the background (black regions in B). D: Laplace filtered image from C. The peaks of the distance transform (the points that are half-way between two boundaries) have lowest value. E: Thresholding the image in D at the 66% of minimal value. Remaining pixels correspond to the peak of the distance transform. F: Projecting the remaining pixels of E onto the orientation of the ellipse (see B), and a nearest-neighbor interpolation of *n_c_* = 7 uniform spaced locations (starting from first to last pixel), results in roughly uniform spaced center line points, c_1_, …, c_7_, (red), which serve as shape descriptors.

To extract this “center line”, we perform a series of transformation, first computing the distance transform *d*(*x*, *y*) of the extracted blob (Fig. 2C). Then we compute the Laplacian of the distance transform (Fig. 2D) and then threshold it at 66% of the minimum, which yields all pixels at the center of the body (Fig. 2E). Then, we project these points onto the major axis of the previously fitted ellipse (see Fig. 2 B, blue line). We then choose *i* = 1,…, *n_c_* equally distributed pixels from those on the center of the body, and call them the center line points **c**^(*i*)^ (see Fig. 2 F, red dots). Finally, from the distance transform, we can estimate the horizontal thickness for each of the center line dots, 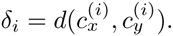) We approximately determine the position of the head by comparing the thickness, since the head region is usually thicker than the tail, in particular for fish (assuming that the video camera is installed from above). If 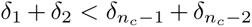 we reverse the order of the center points^11^. Since at least the front part of the body of an animal is usually rather straight and rigid (pointing to the direction of movement), we use the orientation of the connecting line between **c**1 and **c**3 to rotate image patch to a reference position^12^. From this rotated image, we extract an image with fixed dimension to be used as a feature for the identity classification algorithm (see below).

Note that although extracting the center points and thicknesses yields good shape descriptors in particular for fish and are additional useful features for detection-to-track assignments, they are not essential for our tracking approach since it simultaneously relies on other features.

### 2.2 Detection-to-track assignment

Assume that in the current frame *n_q_* objects *q_i_*(*t*), *q_i_*(*t*), *i* = 1,…, *n_q_*, were detected, from which a number of features *f_l_* are extracted, i.e. we write *f_l_*(*q_i_*(*t*)). Given the history of the *n_b_* tracks, *T_k_*(1),…, *T_k_*(*t*−1), for all *k* = 1,…, *n_b_*, which of the detected objects should be assigned to which track?

To solve this assignment problem, i.e. finding suitable *k* and *i*, so that we can safely assign a detection to a track, *T_k_*(*t*):= *q_i_*, we first define a cost function *D_l_* for each of the features *f_l_*. Then the cost of assigning the *i*th detection to track *k* is given by

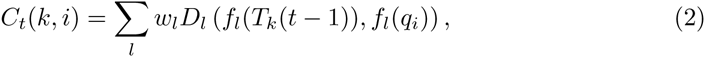

where *w_l_* a weighting factor for individual features. Note that for some tracking methods, new tracks could also be established for unassigned detections, or lost tracks deleted, if assignments could not be made. However, we here consider the number of individuals fixed per experiment and in principle always visible, and thus do not change the number of tracks, once established.

In Eq. 2, we here directly compare a feature of the detection (such as the spatial position) with the feature of the assigned detection of the previous frame. One could also first compute a prediction of the e.g. position, for instance by using Kalman filters, and compare the prediction with the actual location of the detection. However, we found that in case of fish, animals often suddenly changed the direction of movement, so that a linear prediction of the location is often worse than having no prediction at all (given that the frame rate of the camera a reasonable high). In any case, **xyTracker** also supports Kalman filter predictions with a constant velocity model which could be turned on when tracking other animals moving on a straighter path^13^.

In Table 1, we summarized a list of features that could be used by **xyTracker** to compute the cost matrix. The weights *w_l_* in Eq. 2 depend on the history of costs and thus adapt in an online manner to the problem at hand. We set

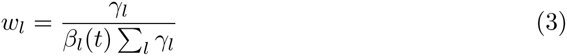

where *γ_l_* are a pre-defined weighting of the diffierent features, defined by the user^14^. The *β_l_*(t) are running means of the distances of individual features, i.e.

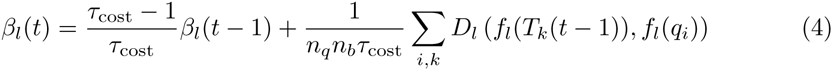

The time constant *τ*_cost_ is set to 500 frames per default. Thus, the absolute values of the distances (which have in principle different units) are automatically adjusted to around 1 for all features^15^.

**Table 1:**
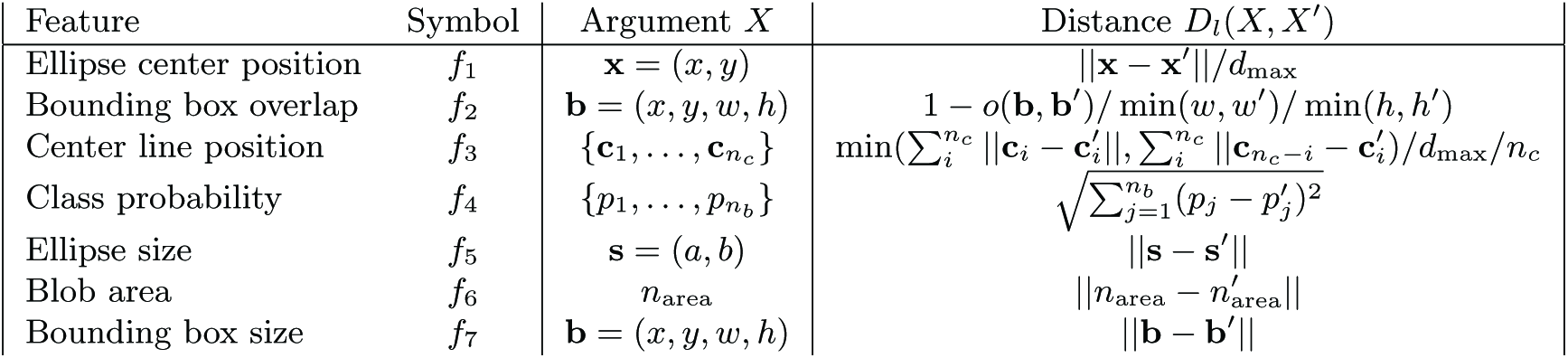
Features and distances used for assigning detections to tracks. In the above, min(*a*, *b*) is equal to a if *a* < *b* otherwise *b*. *d*_max_ = *v*max/*F_R_* is a maximal distance possibly covered by a fish depending on the maximal velocity of the fish *v*_max_ and the frame rate *F_R_*. The function *o*(**b**; **b**’) is the overlapping area of two rectangles **b** and **b**’ (where **b** = (*x*, *y, w, h*) indicates a rectangle with top left corner at (*x*, *y*) and width *w* and height *h*). Further, *n_b_* is the number of fish, and *p_j_* the probability of a detection classified as fish identity *j* (see Methods).

Once the cost matrix *C_t_* is computed, the detections-to-tracks assignment problem is solved by the Hungarian matching algorithm ([32], Munke's variant). However, to avoid wrong assignments of blobs that correspond to noise, we additional include the cost for non-assignments, *c*_non_ (*t*). Setting this value correctly is crucial for good tracking results and we thus let this cost adapt automatically to the problem at hand. We set 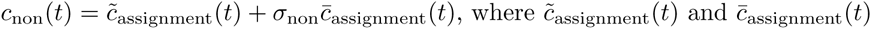 are the running maximal and mean cost of past assignments (with time constant *τ*_cost_, similar to Eq. 4), respectively, thus an average over those elements of the cost matrix *C_t_*_-1_ which correspond to the assigned track-detection pairs (*k*, *i*). *σ*_non_ is a parameter^16^ of **xyTracker**, which can be adjusted if the automatic estimates do not yield in reasonable results. Note that by setting a cost for non-assignments, some tracks might have been lost, that is, they did not get assigned a detection in the previous round. In this case, the features (such as the position) of the non-assigned tracks remain the same in the next frame. Thus, for lost tracks, animals might have moved farther away in the meanwhile, so that a correct assignment generally has increased cost. We thus first handle the matching of detection to visible tracks, as described above. Then, in a second round, we match those left-over detections to tracks that were invisible (that is, non-assigned) in the previous frame^17^. For that, we increase the cost of non-assignment proportional to the number of consecutive invisible frames*, n*_invis_, to 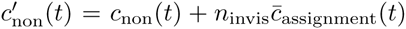. We found that this works well in practice, since it forces assignments for long periods of lost tracks. Note that we additionally test (and correct) for possible wrong assignments with a classifier that learns the identity of each animal (see below). Moreover, we also exclude detections that would overlap too heavily with each other and discard assignments that would result in a movement higher than a predefined maximal velocity.

### 2.3 Recognition of individual animals

The aim of using an identity recognition system in addition to the usual tracking described above is that if two tracks come in close proximity with each other, it is possible that detections are wrongly assigned to tracks, since the cost of assignment becomes similar. Thus, the tracking can be divided into two situations. If animals are far apart from each other^18^, tracking can be done mostly distance-based and is very reliable^19^. However, if two animals come very close and interact with each other, a situation of behavioral relevance, wrong assignment rates increase considerably. Moreover, as pointed out by [45], once tracks are wrongly assigned, the identity of the animals are mixed and errors accumulate, severely compromising the value of the behavioral observation of individuals in groups. In [45], after collecting and tracking the positions, a second pass through the video is done for learning and “sorting” the identity between periods of “crossings”. While an effective approach, we here develop an online tracking system, thus cannot afford to pass through the whole video several times.

Therefore, each frame we test whether two (or more) animals are close to each other, i.e. crossing**. xyTracker** waits for a number of predefined frames after the crossing event to accumulate appearance information, and then judges, based on the accumulated evidences, whether the identities of the tracks after crossing corresponds to those before the crossing event. If a permutation happened, the identities are switched accordingly and trajectories are corrected in retrospect (up to the crossing event). In this approach, although shortly after the crossing the assigned identities might be wrong, the wrong assignments will be corrected on-the-fly, so that errors do not accumulate. In this way, **xyTracker** needs only one pass through the video, so that online-tracking and feedback experiments are possible (see results). We labels this identity handling “switch-based” approach (SWB). Moreover, apart from this explicit switching method, **xyTracker** simultaneously implements a more implicit approach using multiple hypothesis tracking to maintain the correct identity for each track based on unique appearance information of the tracked animals. This directed-acyclic graph-based (DAG) approach is further explained below. Tracking is done simultaneously with both approaches **xyTracker** and tracking results of both approaches can be accessed after tracking is finished^20^.

#### 2.3.1 Features for identity classification

Similar to [45], our tracking system includes a module to recognize individual fish. Thus, we have to extract features to be used for training the fish recognition system (described below), that should be not only unique for each individual animal but also invariant in geometrical transformation. In the situation of a camera viewing a planar scene from above, useful identity features should in particular be invariant to translation and rotation of the body axis.

In the recognition approach by [45], the authors constructed a 2-dimensional histogram of intensity values versus pair-wise distances within the detected blob. Although the authors show reasonable good identification results using this histogram feature, there are several drawbacks with this approach.

First, the histogram computation has to be done for each frame and detected fish and is relatively costly (in the order of *n*^2^, when *n* is the number of pixels of the blob). Second, although invariant against rotation and translation, the histogram is not invariant against object shape deformations. Since e.g. fish bend their tails regularly to move forward, the histogram method thus seems not well suited. Finally, recent success in object classification performance is rooted in the approach to directly take the image intensity values as inputs to a learning algorithms (and learn useful features from the data), without pre-extracting arbitrarily selected features beforehand. It is thus likely that the 2-D histogram neglects some useful information about the identity, which would be present in the raw pictures of the animals.

We here thus simply use the (gray or color) image patch of (the upper portion of) the animal as identity features (see Fig. 3). For that, we use the center line information (or simpler, the orientation of fitted ellipse) to explicitly rotate the extracted detection to an invariant orientation and use the parameters L and W to extract a fixed part^21^ of the (upper) body.

**Figure 3:**
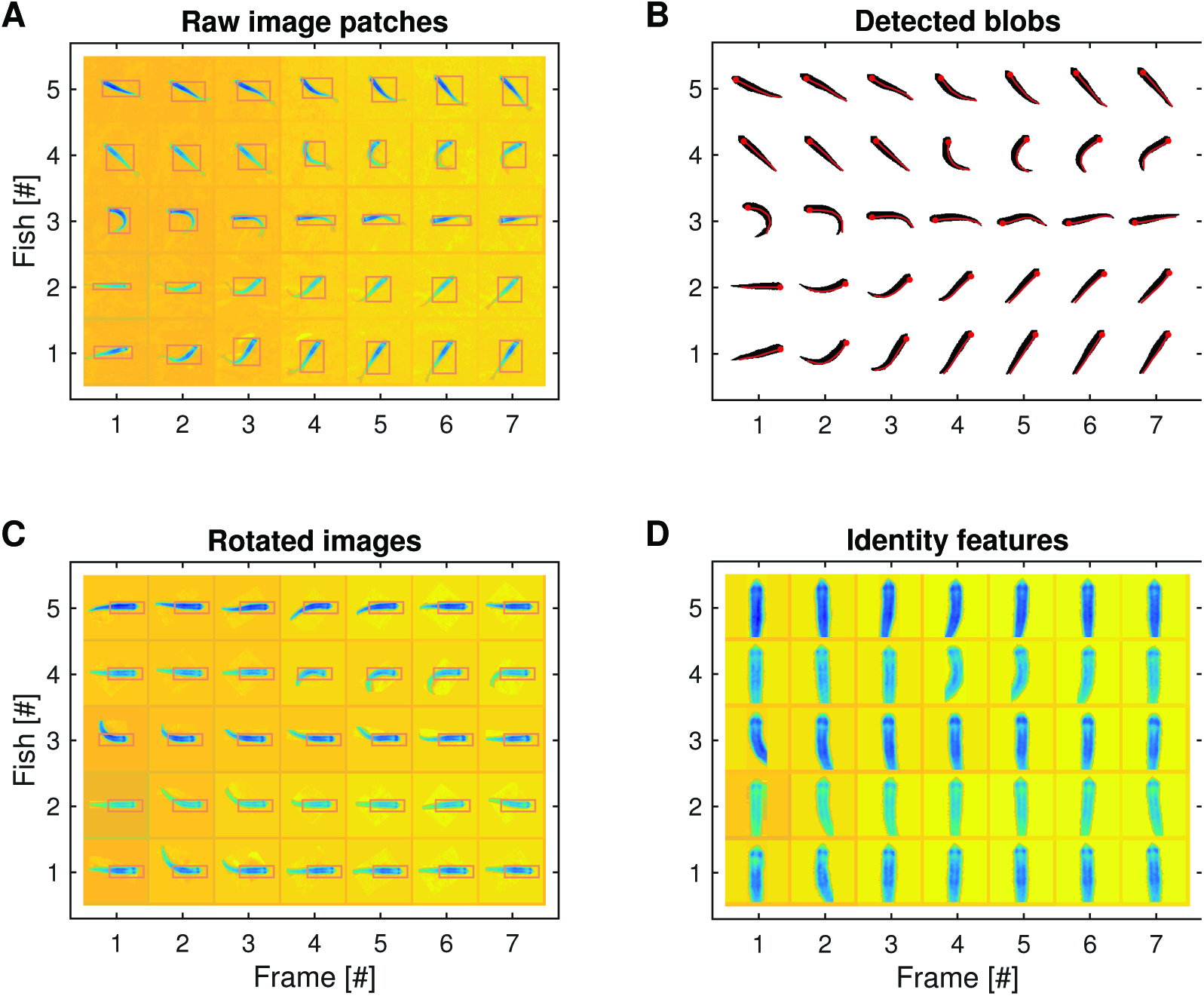
Extracting features for identity recognition of individual animals. A: Raw image patches of 5 individual fish for 7 consecutive frames (example video taken from [45]). Rectangles indicate bounding boxes of detected blobs. B: Detected blobs corresponding to the patches in A. Red lines indicate the center line features from Fig. 2. Red dots show point c1, that is the head position. C: Patches are rotated according to the orientation of the frontal portion of the body centered at the head (red point in B). Rectangle regions indicate the location of the extracted identity feature in D. D: Identity features extracted from C. Note that features are largely invariant in respect to rotation, translation, and bending of the fish.

Taking only the frontal portion in case of fish has the advantage that the movement of the tail becomes irrelevant. For distinguishing the individual fish, it seems that the frontal body part is already very informative (see Results). Note that extracting the identity features in this unsupervised manner allows the identification of any kind of animals. Thus our approach is not restricted to the tracking of fish, similar to the approach by [45].

The image patches are then used by the identity recognition module of the **xyTracker**. Since we explicitly rotate the image patches, our identity features are ensured to be translation and rotation invariant (see Fig. 3D). Thus, we found that a simple classification system is enough for recognition based on this features. Currently, a combination of principle components and Linear Fisher Discriminant analysis is used to reduce the dimensions of the image patches and a Gaussian model for representing the appearance of each identity (see below). However, in principle, a more powerful recognition systems could be used within the **xyTracker** framework^22^.

Note that **xyTracker** assumes that (1) the camera is recording the environment from above, and that (2) the size of the animals does not change severely (i.e. an approximate planar environment, for instance, the water should not be too deep in case of tracking fish). For experimental setups for tracking animals in 2-D, these assumptions are often valid. If the assumptions were not valid, extracted identity features are more variable within an individual but still could be used in our framework. It might, however, require a more powerful (non-linear) classification system, that can handle such variability (see Discussion).

#### 2.3.2 Identity classifier

Our recognition system consists of a classifier, that, when given an identity feature (a local image patch of the detected object, as described above) returns a probability for that feature to belong to one of the *nb* animal identities present in the setup. The classifier maintains internally a model for each identity appearance and is updated continuously to learn a better (or changed) model over time. In principle, the classifier is modular^23^ in xyTracker and could thus be changed or extended if needed.

We found that a simple Gaussian mixture model performed well in practice (see Result section). At the start of the video or recording session, we wait for the first prolonged periods (in practice about 150 frames^24^), where no crossing events were detected^25^. We then use the batch of identity features (see Fig. 3) extracted from each frame to initialize the classifier. The initial batch of identity features per track is used to determine^26^ *n*PCA PCA components on the pooled sample. After dimensional reduction, we further reduce the dimension by selecting *n*LFD components^27^ with largest fisher discriminant between the classes ([39]). In the current implementation, the extracted components are fixed at the beginning for the remainder of the video or recording session and identity features are projected onto the same components. From this projected sample, we compute and update an *n*_LFD_-dimensional Gaussian (sample mean and variances) for each of the *n*_b_ individuals.

#### 2.3.3 Testing the identity

In each frame, for each detection, we compute the identity probability vector **p**_*k*_ = (*p*_1_,…, *p*_*n_b_*_) for each track *k* by calculating the probability according to the Gaussian for each class. Further, we define for each detection a weight *ω_k_*(*t*) according to

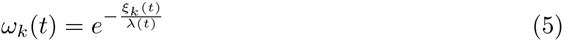

where *ξ_k_* is some positive value indicating the noisiness of the identity feature. We use currently the standard deviation of the center line points, indicating the degree of bending of the body or the deviation from an oval shape and thus a potential noise source. Moreover, we set *ω* to zero if it is smaller than^28^ *e*^−2^/2. We further explicitly set [omega] to zero, when head and tail were mixed up (judged by the velocity direction) and at times when the the assignment cost was too high (with threshold 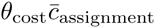). We let 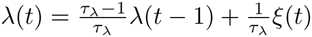, adjust automatically on the time scale^29^ *τ_l_*.

When averaging the prediction over several frames, we use the corresponding weights *ω_k_*(*t*) and compute a weighted mean 〈**p***_k_*〉 of the probability vectors.

#### 2.3.4 Online learning of the classifier

The classifier learns automatically and continuously in on online manner. Similar to [45], we use periods, when no fish is crossing to update the current appearance model in a batch-like fashion simultaneously for all tracks. The adequate number of frames with well separated tracks (“unique” identity frames) needed to trigger the updating depends on the expected average number of track crossings^30^. Moreover, when single tracks are not involved in a crossing over a longer period^31^, appearance features of single identities are independently updated.

During updating, we discard individual samples if they are too noisy (if [omega] = 0). Currently, we use the batch of identity features to update the mean and variance of each Gaussian with time constant^32^ *τ*_class_ (set to 5000 frames by default). Before updating after well separated tracks (“unique” frames update), however, we first use the batch sample to predict the identity for each tracks, yielding a square probability matrix *P*. We use Hungarian matching with cost 1 − *P* to match the samples to the identities. We only update the classifier, if the matching result is consistent with the previous identities of the tracks. If not identical, some undetected mismatching might have occurred during the recent past. If unsure (depending on a forced update probability parameter^33^), the classifier is nevertheless updated ignoring the prediction, or, if sure^34^, the assignment is changed according to the prediction of the classifier and the identity of the tracks is re-assigned in retrospect. When identity switch of the tracks is predicted, this means most likely that an undetected mixing of tracks occurred in the recent past. We search for this switching point from the last crossing point, and look for the points where tracks are either nearby, have a sudden switch in class probability, or are transiently invisible (loosing tracks temporarily is a source of miss-assignments). We use a consensus point between these methods to start the swapping of the identities until the current frame. If more than one track are mixed up (in a permutation of order larger than 2), we use the identities predicted by the DAG approach (see below) rather than simply swapping the identity of two tracks.

#### 2.3.5 Explicit handling of tracks after crossing events

Since most identity switches occur during crossing events, when animals are in close proximity and possibly temporally occluded. **xyTracker** thus includes a system to explicitly tests the identity shortly after a crossing event occurs and correcting the tracks in retrospect, when the identity was indeed switched. Since the identity corresponding to the tracks that crossed are (assumed to be) known before the crossing, only the subset of fish that actually crossed have to be tested for correct assignments. This reduces the number of tests and also the requirements for accuracy: only crossed tracks have to be told apart, not all available tracks

We consider two tracks potentially crossed in frame *t*, if their bounding boxes overlap or their position is closer than the length of the body *L* times a crossing radius scale^35^ λ. A potential crossing is considered a crossing event if either one track is invisible^36^ at time t or the assignment cost of one of the tracks is higher than a threshold 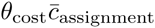. The parameter *θ*_cost_ can be adjusted^37^.

When a subset of tracks crossed, we progress as follows to avoid mislabeling of the identities after crossing (see Fig. 4). Once a crossing event is detected (Fig. 4A, red bounding box), the identity of the tracks which participate in the crossings are remembered. If in the next frames, tracks are still nearby, the crossing zone is enlarged, or, if other tracks get involved, the number of crossed tracks is enlarged (red zone in Fig. 4B). After waiting a fixed number of frames after the last crossing event^38^, the identity features from the tracks involved in a crossing event are taken (light gray zone in Fig. 4B), and tested against all identities involved in the crossing (see Fig. 4C). If *p_i_* are the probabilities if identities would be not permuted and the *p_*σ*_*(*i*) the probabilities of the predicted permutation *σ*(*i*) (using the Hungarian method), we reassign the tracks according to the permutation if the improvement in probability is larger than a threshold, 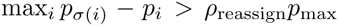, where *p*_max_ is a running average of the maximal classification probability across detections in a frame and *p*_reassign_ a parameter^39^. If class probabilities are consistent with the assumed identity of the tracks for all participating tracks (or below the reassign threshold), the tracks “exit” the crossing event (see Fig. 4D) in case when 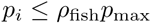 or 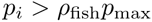 and 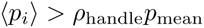 where *p*_mean_ is the running mean of the classification probability across objects in a frame. Otherwise the crossing event is extended into the next frame, since not enough evidence was available (e.g. in this case the grey area in Fig. 4B would be enlarged).

**Figure 4:**
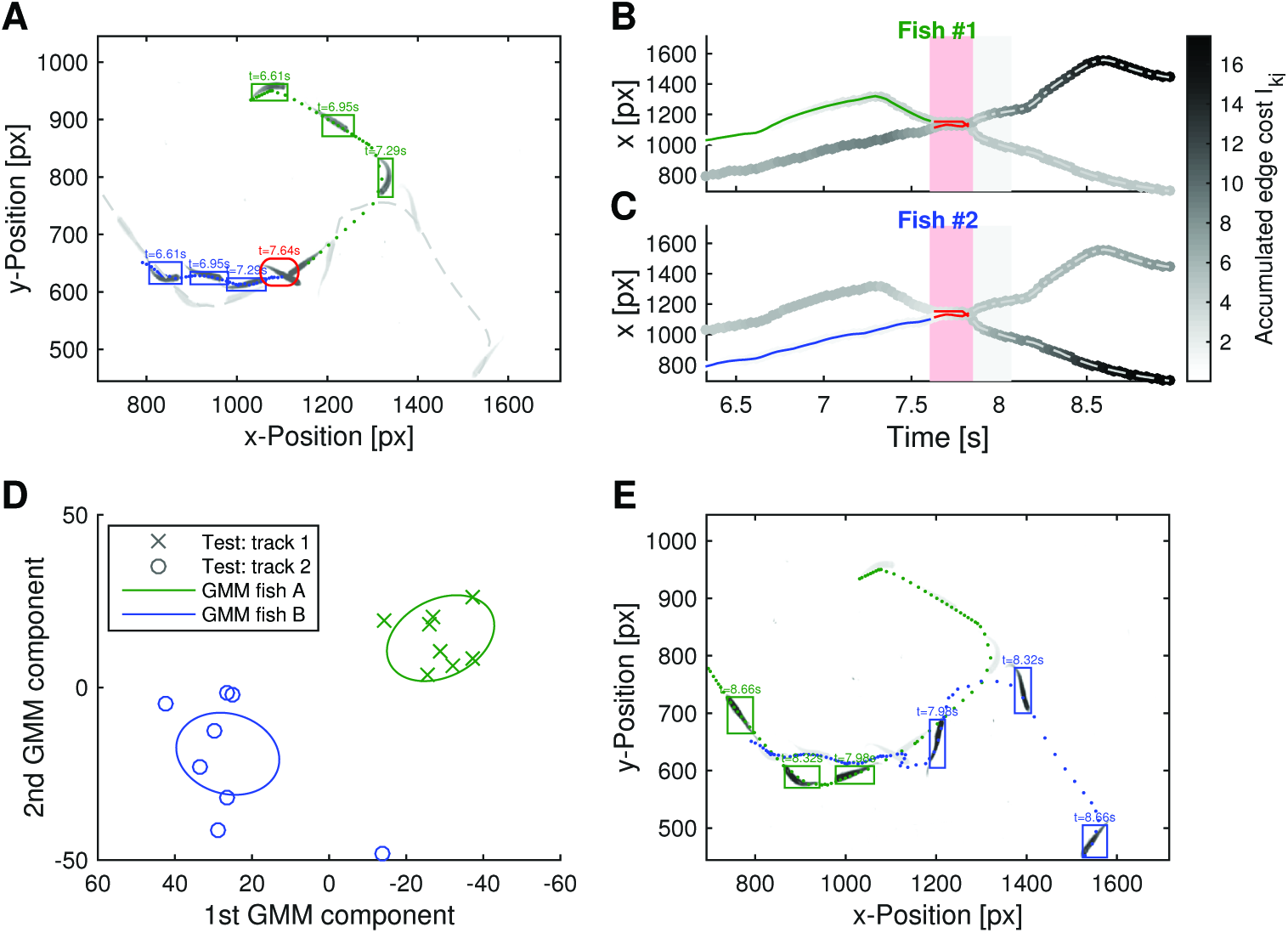
Handling of tracks crossings. A: Two or more animals often interact with each other, coming in close proximity. If tracks are crossing (red) the identity (either blue are green) might be erroneously switched after the crossing (grey tracks). B,C: To prevent this from happening, **xyTracker** uses appearance to predict the identity of assigned objects. The identity features are accumulated based on an online shortest-path algorithm (DAG approach). Note that identity of tracks after crossing can be decided based on the accumulated edge cost (grey level of the tracks). Alternatively, **xyTracker** also explicitly predicts the identities based on features extracted form frames shortly after the crossing (gray area; SWB approach). D: Appearance is compared to the learned appearance model. Identity feature are tested against the Gaussian mixture model (GMM) for each animal participating in the crossing (note that the GMM is learned online at times when no crossing is detected). E: In this manner, the identity of each track can be established and the tracks are re-assigned in retrospect if switching happened during crossing.

If a switching occurred after crossing, identities of the tracks are permuted and trajectories are adjusted in retrospect up to a consensus point in history, analogous to the reassignment before updating the classifier as described above. Thus, in case the identities of the tracks need to be reassigned, note that **xyTracker** reports the wrong identity until it switched the fish tracks. While this is of no consequences for offline video tracking, it might compromise the stimulus presentation during online closed-loop experiments, since the identity information is reported wrongly for a short while. However, in practice, if animals are not too heavily interacting, this time of uncertainty is rather short (e.g. about 5 frames after a crossing) and can be adjusted^40^. In any case, since the identity of the tracks will be corrected, identity errors do not accumulate.

#### 2.3.6 Identity tracking by the calculating the shortest-path on the directed-acyclic track hypothesis graph

A powerful and popular method for tracking multiple objects in computer vision is the multiple hypothesis tracking (MHT) approach [47], which can be combined well with appearance information [30], like our identity features. In this approach, multiple hypothesis of tracks are kept simultaneously and an optimization is applied to find the most probable tracks per identity afterwards (see also review [12]). Thus, since multiple hypothesis about possible trajectories are kept simultaneously and disentangled over time, this approach is more powerful and general than the described explicit switching approach above. A problem of the original MHT approach is, however, that the number of possible hypothesis grows exponentially in time and thus unlikely hypotheses need to be discarded during tracking [20, 12, 30]. In track-oriented MHT, each track (aka individual animal in our setup) is maintained per frame [12]. Since for the tracking of animals in 2D in an experimental setup, the number of animals tracked is typically small (below 100) track-oriented MHT is nevertheless very efficient in our implementation.

Related to the approaches of [3, 46, 56], we developed a fast greedy online version for tracked-oriented MHT to handle the track identities in combination of the online-learned appearance model. Here, we use a number of simplifying assumption. First, we can assume that the number of animals in the experimental setup remains the same throughout the experiments (although individual animals are allowed to be “lost” for a number of frames). Second, our appearance model is dense, that is, in each frame a noisy but relative informative identity feature is extracted. This is achieved by extracting identity features as described above. Further, we combine the MHT approach with the detection-based approach described above. That is, only assigned detections per frame, or predicted positions of lost tracks, are included in the MHT approach. We thus follow a detect-then-track approach for determining the identities with our MHT method [56]. We proceed as follows.

Let *q_j_*(*t*) be the *n_b_* detections that would be assigned to existing tracks as described above. We use these to form *n_b_* graphs of *n_b_* hypotheses, one graph for each identity *k*, in a similar manner as proposed by [46]. In the frame *t*, assume for the track hypothesis *T_ki_* that the assigned detection in step *t* −1 would be *T_ki_*(*t* − 1):= *q_i_*(*t* − 1). Then all possible *n_b_* new hypothesis involving this track would be formed by assuming any of the current *n_b_* detections *q_j_*(*t*) would extend the existing track and form a new one of length *t*. When defining a distance (or likelihood) between the detection at *t* − 1 and those at *t*, i.e. 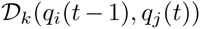, that can be attached to an edge, the track hypotheses form a directed graph which number of nodes grow exponentially in *t*. Finding the best hypothesis can then be cast into a constrained optimization problem finding the path with least cost, the shortest path [18]. The main constraint in this optimization problem is usually that each detection has to be uniquely assigned to one track for each frame, resulting in a linear programming problem [3, 46].

However, we found that simply computing the shortest-path for each identity individually, without constraint, already yields very good results in practice and can be done naturally in a very fast online manner (without the need for overlapping windows with limited time horizon for the linear programming approach [46]). This is because only a small number of fish are crossing simultaneously at a given time while others are spatially well separated, and because dense appearance information is available using our identity feature extraction.

We derive our method by realizing that the hypothesis graph for each identity *k* is a directed acyclic graph (DAG) in topological sorted order (see [18] for details on the theory of graphs; see also Fig. 5). Moreover, since there are no edges between nodes within a time frame, the value for a possible shortest path involving a particular detection can be calculated for each detection in a frame independently (see Fig. 5 for an illustration). Therefore, if assumed that the number of detections and identities per frame are fixed to *n_b_*, computing the shortest path reduces to updating each *t* a cumulative cost matrix *I_kj_*. We need to evaluate for each of the *n_b_* possible shortest paths for identity *k* of length *t* − 1, with *T_ki_*(*t* − 1) = *q_i_*(*t* − 1)

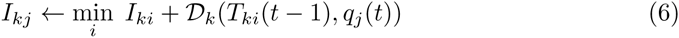

and save the selected *i*\* for each *T_kj_*(*t*). Intuitively, the shortest-path involving the current detection *q_j_*(*t*) is that path, which edge transition cost from any of the previous detections *q_i_*(*t* − 1), plus the accumulated previous edge transition costs until arriving at *q_i_*(*t* − 1), is minimal [18]. In this manner, each time step *t*, exactly *n*^2^*_b_* shortest-path hypothesis are computed and kept, that is, *n_b_* shortest paths up to *t* for each of the *n_b_* identities.

**Figure 5:**
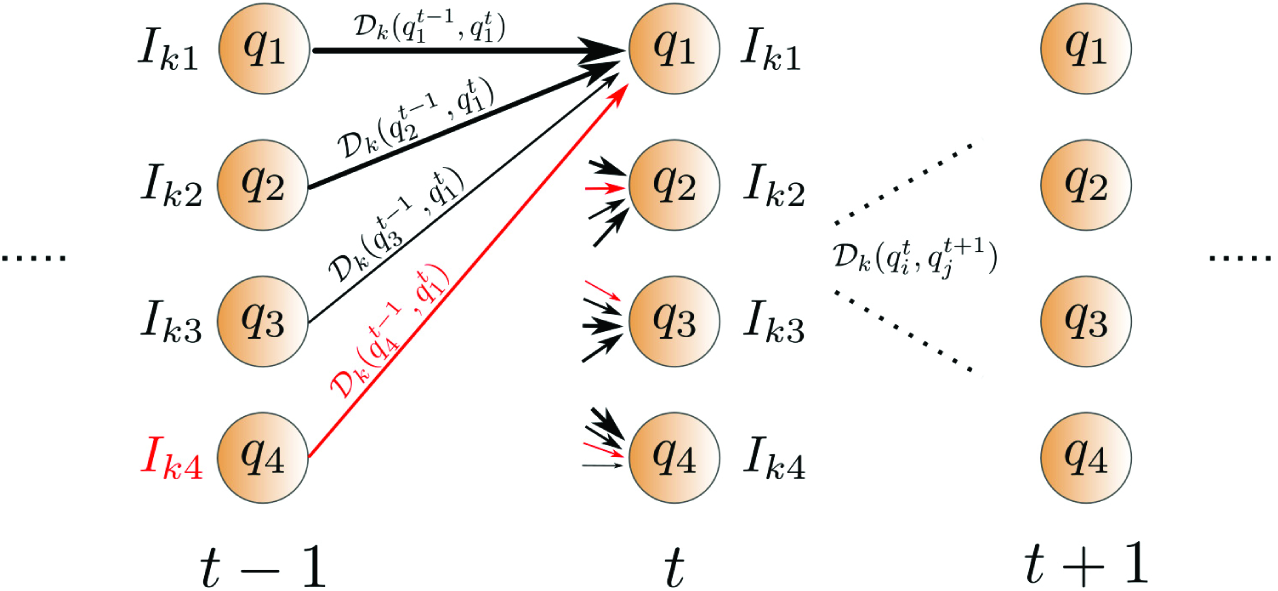
Multiple hypotheses tracking (MHT) using online shortest-path computation. For each identity *k* a directed-acyclic graph is built. Each node represents a detection which was assigned to an existing track. Cost of edge transition is given by the cost 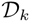, which includes a term for spatial distance and the appearance determined by the classifier from the identity feature (see Eq. 7). The shortest-path is computed by updating the cumulative cost matrix *Ikj* and choosing the minimal path (red) involving detection *q_j_*(*t*) according to Eq. 6 (in the scheme shown only for *q_1_*(*t*)). The shortest-path involving a given node at *t* can be back-traced (by following the red arrows backwards).

Note that for each of the *k* identities, our algorithm computes the globally shortest path in respect to the edge cost. However, in principle, parts of the tracks for two identities might be overlapping, that is, some detections might be assigned simultaneously to multiple identities. To avoid that a particular identity follows long stretches of a track of another, we let the cost matrix 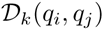 (governing the edge cost) depend on the appearance of the identity *k*:

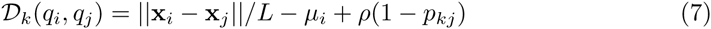

where *p_kj_* is the probability of the detection *q_j_* belonging to the identity class *k*, determined by the classifier and the identity feature described above. The weight^41^ p varies the inuence of the accumulated spatial distance relative to the accumulated identity probability on the edge cost. The term *µ_i_* is the minimum spatial distance between the previous and all current detections, 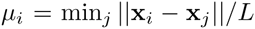 so that a distance penalty is added to the cost, if not the nearest detection is taken. When tracks are “lost”, that is, no matched detection can be found matching to a track at time *t*, we simply reuse the detections from the previous time step^42^. Note that Eq. 6, by subtracting the minimal distance, mainly accumulates the class probability for a particular identity on a track. When tracks are nearby, hypotheses follow both tracks until the next crossing point, where only the path along which the appearance cues did more likely resembled the identity will win and the other track be forgotten.

The globally shortest-path for each identity is only determined when tracking is finished. To access the current estimate of the identities for each detection, to be used for online stimulus presentation (see below), we compute a Hungarian assignment on the matrix *I_kl_*, which yields a unique assignment per individual (without overlapping). Note that we initialize *I_kl_* at the beginning with 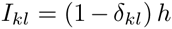, where *h* is a high cost value, to ensure that the starting node for each shortest path is from unique individuals. The end-node for each track is determined by the identity prediction from the Hungarian method. In the implementation, the positions for the globally shortest paths are in retrospect determined by back-tracing the references for each edges. Therefore, our DAG method is very fast, since it computes the globally optimal shortest-paths in 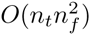 time.

In **xyTracker** both, the switch-based (SWB) and the DAG-based tracking are computed simultaneously^43^ and resulting tracks can be chosen after tracking has finished for further analysis^44^. In practice, both methods are advantageous over the other for some applications. In general, if appearance features are very informative about the identity the DAG approach is advantageous. This is mainly because the consensus crossing points is difficult to be determined in retrospect for the SWB approach, in particular, when more than 2 tracks are crossing multiple times. Then there might by multiple instances of track switching^45^.

If appearance cues are less informative and/or tracks are crossing often, for instance when many small animals (e.g. > 20) are tracked simultaneously, the tracking is generally more challenging. In this case, the DAG approach might result in many overlapping trajectories, and thus the SWB approach should be preferred, since it yields more stable identities for each track.

In case when appearance is well discernible, we found that not having unique detections to track assignments in the DAG approach is in fact advantageous in some cases. For instance, during occlusion in crossing events, one detection indeed belongs to two tracks rather than one (e.g. fish shapes overlap and only one blob with the shape of a cross is detected). Long overlapping sequences are avoided in practice by accumulating the probability of the appearance along the path, making long overlapping tracks unlikely.

For closed-loop experiments, the DAG approach with added Hungarian switches identities too often shortly after crossing events. This might be problematic for some experiments and thus we chose^46^ the more stable identities and occasionally switches of the SWB approach for our closed-loop experiments (see Results).

### 2.4 Stimulus presentation

Since **xyTracker** tracks in real-time in an online manner, it can be used for virtual reality experiments. In this case, the video frame is grabbed directly from the camera and immediately processed by **xyTracker**. After a frame is processed (object detection and assignments to tracks), a (visual) stimulus presenter is called to update a visual stimulation screen (or any other type of stimuli such as electric shocks). **xyTracker** connects to the PsychToolbox in Matlab for stimulus presentation (with OpenGL), a toolbox which is widely used in cognitive research [31]. The base stimulus presenter can be overloaded and extended to easily implement any types of stimulation protocol^47^.

Note that the stimulus can depend on the actual position, speed, direction of movement or any other extracted feature of the collective motion of the group, opening up new directions for studying group behavior (see Results for an example closed-loop experiment).

### 2.5 Implementation

**xyTracker** is freely available for academic research^48^. **xyTracker** is implemented with Mat-lab (2014b, MathWorks, Natick, USA) relying on and C++-core (using the OpenCV library^49^) for performance reasons. Since **xyTracker** is object-based, it is thus modular and easily extensible. We implemented three different versions depending on the available libraries and operating system^50^: (1) a platform independent version using only Matlab functions (and toolboxes) (2) (some what) platform independent using Mex-OpenCV interface^51^ (3) on a Linux system using performance enhanced multi-threading C++/Mex core which interfaces to a Matlab front-end. Performance and feature set^52^ of the three versions differ. For tracking 5 medaka fish for the initial 100 seconds of a reference video (available from [45]) the different versions needed 217, 110, and 46 seconds^53^, respectively, on the same quad-core personal computer (HP Z230). Note that version 3 tracks significantly below real-time (100 s) and should thus be used for all performance relevant applications. For the complete ca. 8 min video, xyTracker needed about 3 min and 25 seconds to track the fish identities (the purely Matlab-based version 1 is expected to be about 4-5 times slower). However, even the slowest version of **xyTracker** yields a large speed-up in comparison to available tracking solutions with identity detection. For instance, the idTracker software [45] needed for the same video on the same computer 1 h 54 min 36 s (including 4 min 26 s start-up time, where the video was loaded into memory; employing distributed computing with 4 Matlab workers, one for each core) with high memory usage, since the whole video needed to fit into memory. Thus, **xyTracker** delivers a > 30× run-time speed-up to comparable tracking software without significant loss of accuracy (see Results).

Moreover, **xyTracker**’s memory usage is low, since the frames are read in sequential manner and identity features for individual animals are only stored as long as needed (they are removed from memory when the classifier was updated). For example, we tested it on a > 8h color video with about 3 megapixels per frame and color with 40 Hz frame rate (1196000 frames; compressed video file size 6 GB, about 9 TB if uncompressed) without encountering any memory issue on a personal computer (see Results; Fig. 7). It took around 7.5 h to track this video of 3 zebra fish with **xyTracker** on the same personal computer as above. Note that such a long video would have been impossible to directly track with the **idTracker** software in reasonable time^54^.

Furthermore, **xyTracker** supports grabbing and simultaneous saving (and background video encoding with h264/5 codecs) from a video camera^55^. Up to now, xyTracker only supports grabbing from PtGrey cameras via the FlyCapture2 SDK^56^, although adding support for other cameras through OpenCV's VideoCapture functionality would be straightforward. Video encoding is based on the multi-threaded FFMPEG library^57^ and done on separate threads and thus, given enough available cores, does not interfere with the tracking process (we tested it on a HP Z820 Workstation with 16 cores (dual Xeon(R) CPU E5-2690) and found that tracking and encoding can be done simultaneously in real-time). While grabbing, the **xyTracker** will skip frames automatically and thus works on a variable frame rate, if tracking cannot be fast enough to perform in real-time on all frames^58^. Therefore, the stimulus presentation will not lag more than the total time to process a single frame.

### 2.6 Matlab interface usage

For installation information, we refer to the code documentation^59^. **xyTracker** includes a simple run test for checking the code integrity^60^. After successful installation and compilation the test can be called from the command window^61^ by typing

~~~
>> xy.Tracker.runTest()
~~~

For each application (tracked video file or real-time experiment) a new instance of a **xyTracker** object has to be generated, e.g. by typing (for tracking the video **vidFile** with 5 animals)

~~~
>> T = xy.Tracker(vidFile,'nindiv',5);
~~~

A number of options can be given, e.g. in the following way

~~~
>> opts.blob.colorfeature = true;
>> opts.useMex = 1;
>> T = xy.Tracker(vidFile,'nindiv',5,opts)
~~~

Note that options can be given^62^ either as “**optionName,value**” pair or as a structure, or both^63^. For a list of possible options and documentation one can simply call (without arguments)

~~~
>> xy.Tracker()
~~~

After instantiating a **`xyTracker`** object, **`T`** in the above commands, tracking and displaying can be done by a number of methods. For instance,

~~~
>>T.setDisplay(1)
>> T.track();
>> T.save();
>> T.plot();
>> T.playVideo();
~~~

will start to track the whole video file (with online progress display), save the whole object **`T`** to a file (for later reference) and **`T.plot()`** will plot the traces and some analysis results, like a probability map. **`T.playVideo()`** overlays the tracking results with the video for verification.

After tracking, the resulting traces and features can be further analyzed. The results structure will have fields for each feature (i.e. location and orientation) and can be access by

~~~
>> res = T.getTrackingResults();
>> plot(res.t,res.tracks.location(:,:,1)) % plots the x-locations over time
~~~

Please consult the documentation and the code for more information.

### 2.7 Experimental setup

Zebra fish used in our behavioral study were obtained from a local pet shop and were kept together in a filled 45 cm cubic glass tank with heater (around 22◦C) and water filter (Tetra, Spectrum Brands, Madison, US). We custom built two setup for video recording. The first, without visual stimulus, was used for open water tasks. It consists of a 45 cm cubic glass tank, fitted with non-transparent greenish plates to avoid reflections (FR-4). For video recording, we used a Point Grey Grasshopper camera (GS3-U3-41C6C, Richmond, Canada) with high speed CMOSIS (Antwerp, Belgium) CMV4000 2048 × 2048 pixel 1 inch color CMOS sensor connected via USB3 to a HP Z820 Workstation (dual Intel Xeon CPU E5-2690, 2 × 8 cores, 128 GB RAM memory) running Linux (Fedora 21). We used a 16 mm lens (Shenzhong Zhongxin Technology, Shenzhen, China) installed about 70 cm above the water level and with an attached CPL filter (NiSi, Zhuhai, China) to reduce water reflection. Ambient lightening was provided by LED lights.

The second setup was used for visual stimulation. Note that for visual stimulus presentation from below it is necessary to track animals using infra-red cameras and appropriate visible light cut-off filter to avoid that the visual stimulus impairs the tracking. The stimulation setup consisted of an acrylic water tank (28×36 cm, height 20 cm) which was placed on a 17 inch TFT monitor (M-PJ-170, Yinghao Electric Technology, Zhuhai, China, industrial housing, 4:3, 1280 × 1024, 400 cd/m2). The monitor covered the whole tank except for a small border (less than 1 cm). The inner sides of the tank were fitted with matte semi-transparent foil to reduce reflections. For recording with this setup, we used another Point Grey Grasshopper camera with 1/1.8 inch CCD (ICX687, Sony, Japan) and 1928 × 1448 resolution (GS3-U3-28S4C). We removed the IR-cutoff filter of the camera and fitted an IR-pass filter (720 nm, GreenL, Shangyu, China) and a CPL filter (NiSi, Zhuhai, China) on a 12 mm lens. The distance to the water level was similarly about 70 cm. We used 3 custom-made 850 nm LED-light machine vision bulbs for illumination (MVIR0460, Herowei, Hongkong, China).

Visual stimulation were delivered using the PsychToolbox for Matlab by custom written scripts during tracking with **xyTracker**.

### 2.8 Comparison of classification methods

For validation whether identity information is present in our identity features and to compare our simple online classification method (GMM) to state-of-the-art classifiers, we performed the following comparisons.

Using the example video of [45] as a benchmark, we extracted the identity features of the 5 fish for each frame. We validated that **xyTracker** performed similar to the our ground truth provided by the **idTracker** method (see Results). We extracted image patches for individual fish (identity features) by using **xyTracker** as described above. After discarding frames where a fish were not successfully detected, the data set consisted of *N* = 12042 frames with 5 small image patches of size 19×57 pixels showing the individual fish. For training a classifier the first 10000 frames were used as train set, the rest for testing. Before separating the test and train set, we did not randomize the order of the frames to avoid that a classifier could exploit temporally correlation in the feature appearance (feature in nearby frames typically look very similar due to the high frame rate). We used linear classification (linear regression plus threshold activation), logistic regression, multi-linear perceptron (MLP) (e.g. [11] for a review), and convolutional networks (CNN) [34] for classification of the identity of the 5 fish. Pixel values were rescaled to the range 0 to 1 and mean subtracted. We use the python toolbox THEANO [9] to implement the training.

The MLP model had two hidden layers (800 and 500 units, respectively) and one readout layer. The 5 class labels were obtained by soft maximization of the read out activations. Following [43], we use threshold-linear activation functions. The CNN is similar to [34] and consisted of two convolution-pooling layers (20 and 50 kernels, respectively, kernel size 5×5, pooling size 2×2) followed by two hidden layers (800 and 300 units, respectively). In contrast to [34], the convolution layers were fully connected. To increase the competition between features in the convolutional layers, we normalized each pixel in the feature maps by the activations of the same pixel location in the 5 adjacent feature maps [28]. We used a threshold linear activation function and adaptive gradient [23] to optimize back-propagation to learn the weights (batch size is 100, initial learning rate 0.001, decreasing to 0.0001, maximum epochs 200). Computations were performed on a NVIDIA Tesla K40 m GPU and took about 3 min, 10 min, and 20 min to train logistic regression, MLP, and CNN, respectively.

## 3 Results

We developed a new tracking system, **xyTracker**, for monitoring group behavior in a 2D environment. Our approach comprises a fast online tracking method together with an identity recognition system (see Methods for a detailed description). For classification of the identities, local image patches of animals are extracted and explicitly rotated according to the main body axis (thus ensuring translation and rotation invariance of the features), so that simple classifiers can be used to learn the appearance of each individual in an online manner to distinguish highly similar individuals in the group. Thus our tracking platform is not only fast (> 30× speedup to comparable software), it also ensures that identities of the tracked animals are not mixed up over time. Moreover, because tracking and identity learning is done on-line and in real-time, stimulus can be presented in closed-loop experiments based on the current position of any of the tracked animals.

In the following, we first validate our software against a comparable approach, and then perform a number of experiments to exemplify the long-term stability of the tracking, the visual stimulation capabilities, and a simple closed-loop experiment.

### 3.1 Validation of tracking method

We validated our **xyTracker** software with an available tracking method that also accounts for the identity of individual animals [45] (in the following called **idTracker**). For a fair comparison, we use the video of 5 medaka fish provided with the publication of [45] for comparison of our approach^64^. Using this video as a benchmark, we tracked the fish with both methods, and compared the distance of the resulting tracks. Tracking the 5 fish in the ca. 8 min long video with **xyTracker** was completed in about 3.5 minutes on a personal computer. On the other hand, tracking the same video with **idTracker** took almost 2 h. **xyTracker** thus was about > 30× faster than **idTracker** (see Methods for details). Note that for both methods the naming of the fish were arbitrary. We thus adjusted the naming of the fish between both methods by matching the trajectories minimizing the average distance across the video.

We found that **xyTracker** accurately tracked fish by identity. In Fig. 6A the mean distance of the tracks of the 5 fish between the two methods are plotted. Note that the distance is very small for most of the time (considerably below 1 body length), indicating that both methods extracted the the same tracks with the same identity. Assuming that idTracker found the correct tracks without switching (as was indeed confirmed by hand in [45] for this video file), our validation suggests that **xyTracker** successful tracks fish by identity. We found that the frequency of temporary lost tracks is very similar between the two methods (see Fig. 6B) indicating similar detection quality on this video.

**Figure 6:**
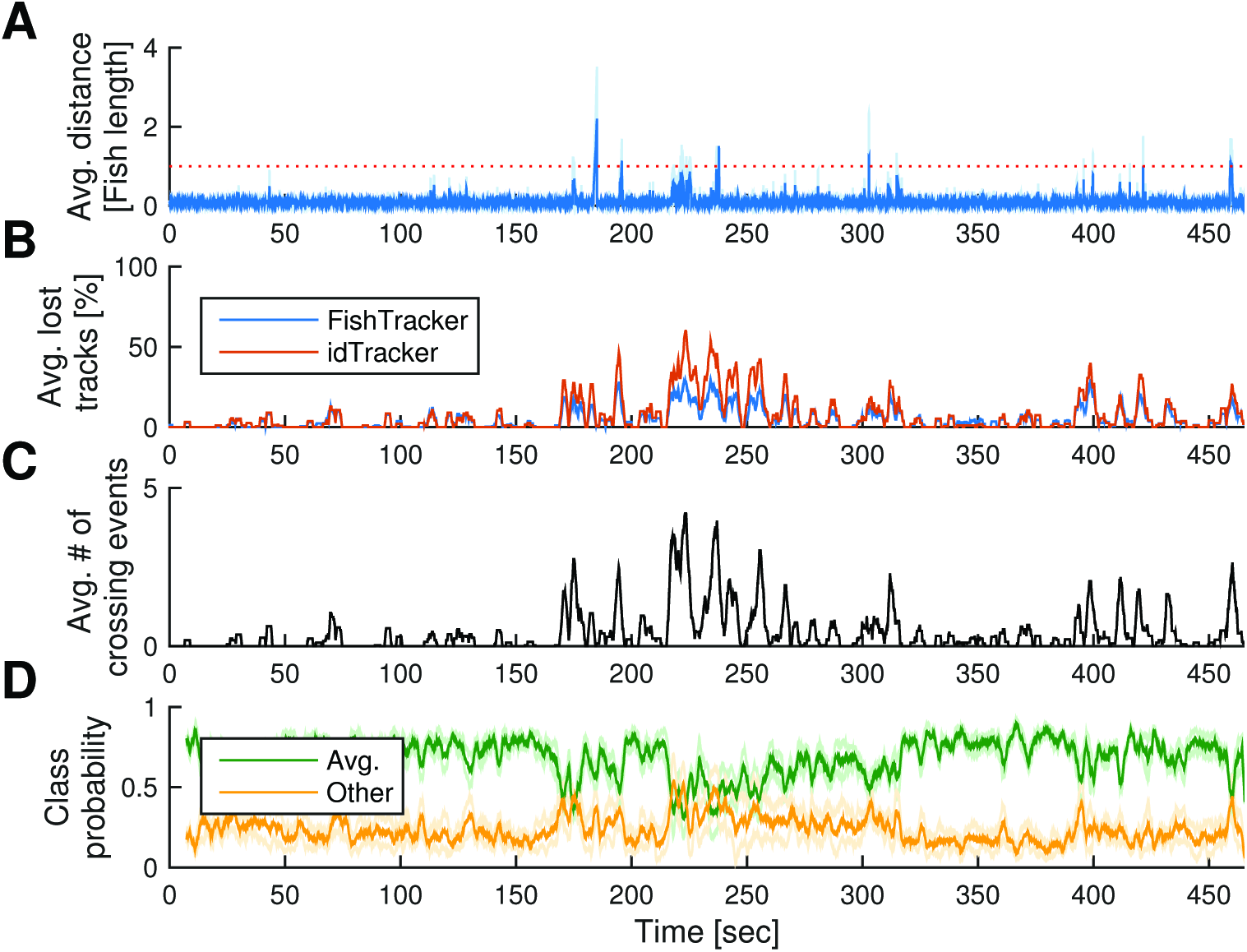
Validation of the accuracy xyTracker with idTracker software. A: Average distance between resulting tracks of the two methods. Note that occasional difference get corrected immediately. Note further that distance reduce to nearly zero after a misalignment due to heavy crossing in the region of 200 s-250 s. B: Detection quality of both methods are very similar. Both methods cannot detect blobs in particular when fish are crossing (they form a single blob instead of two). C: Number of fish crossing, according to the **xyTracker**. Note that many fish are mutually crossing in the 200 s-250 s region, challenging both tracking methods. D: Evaluation of the performance of the identity classifier. Average probability of the selected identity for each track (green line) is compared to the maximal probability when another identity would have been selected instead (“Other”, yellow line). The more both probabilities differ, the more specific the appearance model is learned and the high the confidence in the assignments of the identities to the tracks.

The main advantage of **idTracker** over straight-forward tracking methods, is that occasional mixing of identities do not accumulate over time. This is also true for **xyTracker**. In the video, at around 200 to 250 seconds, the frequency of crossing of fish is very high (see Fig. 6C). During the same time, tracks of both methods differ occasionally(compare to Fig. 6A), possibly because many tracks are lost during this time and wrongly reassigned (compare to Fig. 6B). However, for the remaining part of the video, both methods again yield identical tracks (i.e. the average distance is very low, Fig. 6A), indicating that identity information was successfully recovered by **xyTracker**.

Our method provides a probability for each frame that the currently detected animal belongs to the identity class of the track. Comparing this probability with the “other” probabilities, e.g. by computing the maximal probability when the current animal would be assigned to a different identity, is a good measure of how specific and accurate the identity classifier performs. In Fig. 6D the running average of both probabilities is plotted. The difference between both probabilities is high for the majority of the time, indicating that the individual fish could be distinguished well based on appearance alone. Moreover, during the heavy crossing period, the difference reduces, indicating that during this time the assignment of track identities has lower confidence.

### 3.2 Validation of the identity features

In **xyTracker**, identity is tested using appearance features extracted from the individual animals during tracking. In [45], authors proposed to used a 2D-histogram, which is invariant against rotation and translation of the animals. Rotational invariance is in particular crucial for setups where the camera is mounted from above and animals are free to move in every direction on a planar environment, such as fish in a tank of shallow water. We here propose to use directly the detected image content of each animal instead of computing a histogram of the detections. To ensure rotational invariance, necessary for a simple classifier to recognize individual animals accurately, we simply rotate the detected images along their elongated body axis, which is extracted for each animal during tracking (Fig. 3; see Methods for details). In the following we asked, how much information could be extracted from these features, if one would be free to choose any powerful classifier? Is there enough information to classifier the identities purely based on this feature for any moment in time?

To test how accurately individual fish can be recognized in the example video (from [45] as in Fig. 6) by using solely the identity features, we first extracted the identity features from the 5 fish and labeled the images based on the detected identities with **xyTracker**. We then tested whether the five identity classes could be labeled correctly by using conventional supervised machine learning approaches (see Methods for details).

We found that classification of the 5 fish using the identity feature could be done with high accuracy (> 80%) even using simple methods like linear regression when they are rotated according to the body axis (see Table 2). State-of-the art methods, like convolutional networks [34], reached an accuracy of 97:1%. This result shows that the extracted identity features indeed provide enough information about the individual identities of the fish. We found further that when images patches are not rotated to a reference orientation, simple classification methods, including the GMM approach of **xyTracker** performed poorly (see Table 2; “not rotated”). Only the powerful CNN method was able to learn the rotation invariance from the data.

**Table 2:** Supervised classification of identity features. Fraction of correct classification of the 5 fish for different methods. When the image patches are explicitly rotated to a reference orientation (based on the shape of the detected blob; as described in Fig. 3), even very simply classifiers, such as Gaussian mixture models (GMM), performs well. However, if the raw patches are used (not explicitly rotated; “not rotated”), only the powerful CNN methods learns the rotation invariance from the data.

| Method | Image feature type | Accuracy [%] |
| --- | --- | --- |
| GMM | rotated | 82.0 |
| GMM (5 frames) | rotated | 89.9 |
| Linear regression | rotated | 84.5 |
| Logistic regression | rotated | 88.0 |
| Multilinear perceptron | rotated | 92.0 |
| Convolutional network | rotated | **97.1** |
| Logistic regression | not rotated | 52.6 |
| Multilinear perceptron | not rotated | 58.3 |
| Convolutional network | not rotated | **94.6** |

**xyTracker** uses a simple online learning scheme to form an appearance model for each animal identity. In particular, we fit a Gaussian probability distribution to a number of components of the identity feature (see Methods for details). This simple model has the advantage to be very fast and easy to train (only the mean and variance of a Gaussian per identity needs to be updated). Surprisingly, this simple method already reaches 82% accuracy on the test set, indicating that, when image patches are oriented to a reference orientation, this simple models is a good compromise between accuracy and speed. In **xyTracker**, the identity classification method is used to ensure tracks’ correct identity after crossing events, where the majority of wrong assignments and mixing of identities occurs. Moreover, **xyTracker** combines identity features of a number of consecutive frames to yield an further improved accuracy. For instance, when combining estimation of 5 consecutive frames, the accuracy of the GMM method increases to 89:9% (see Table 2). Since **xyTracker** usually accumulates evidence along an even longer tracked path, multiple frames will be combined before critical decisions at crossing points of two tracks have to be done. We thus conclude that the identity feature and classification method used in **xyTracker** is appropriate.

### 3.3 Territorial domains

We observed that zebra fish, given visual reference cues in the aquarium such as stones, tend to establish their own territorial domain after several hours, as was reported previously [51, 53, 52, 45]. Investigating this behavior is a good practical benchmark for a tracking software like **xyTracker**, since the individual identities of the fish have to be maintained over a long period. If the resulting trajectories would not show such a segregation of identities in space, it would mean that identities were not maintained and an accumulation of track mismatch errors occurred over time.

We thus recorded a > 8h video (having in total 1196000 color frames with a resolution of 1778×1760 pixels) of three behaving adult zebra fish (see Fig. 7A). To establish a visual reference for domain behavior, we placed a number of small stones in the middle of the aquarium^65^. Tracking the zebra fish in the recorded video is challenging for a number of reasons: (a) The video size is very large, about 6 GB compressed (9 TB uncompressed), thus requiring low memory usage. (b) The recorded time is long, thus the identity of animals has to be maintained for a long period of time. (c) Fish are small compared to the environment and relative contrast against the background is relatively low. (d) The background is cluttered with small stones, where the color of the stones is similar to the color of the fish. (e) The background changes over time because the stones tend to move, caused by the movement of the fish. (f) In this video, fish tend to interact heavily with each other, with fast bouts of swimming, driving each other out of their own territory (example of a short hunt between fish 2 and 3 is shown in Fig. 7 A). Thus, a further challenge is that such dynamic interactions between fish tend to disturb the surface of the water, which momentarily changes the background and thus causes a number of wrong blob detection that should be ignored by the tracking software or otherwise may lead to lost tracks (see Methods for a detail description of the tracking process).

**Figure 7:**
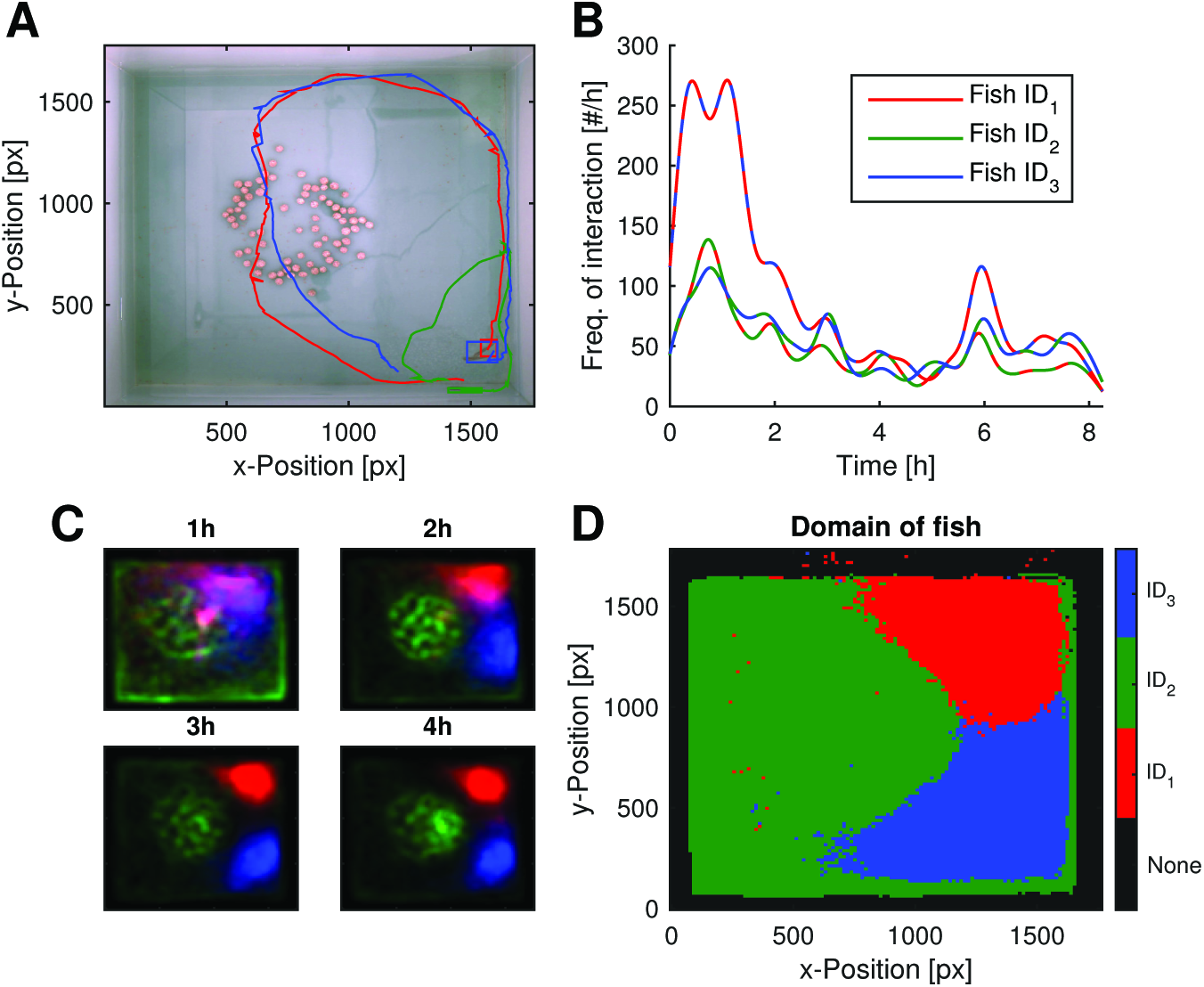
Emergence of territorial behavior. A: Environment used for this experiment. Some movable round stones are placed in the middle of the tank. Three zebra fish (rectangular bounding box) interact frequently showing aggressive and fast escape behavior (tracks for a few seconds are shown in solid lines). B: Frequency of high velocity bursts (> 20 pixels/frame) of at least 3 consecutive frames, in which two fish participate. Colors of fish identities as in A, dashed lines of two colors indicate interacting fish according to the legend. Note that the two smaller fish, 1 and 3, interact much more often at the beginning presumably to fight for territory. C: Emergence of domains. Accumulated locations for each fish in false color. Whitish regions indicate regions that were visited equally often by all fish. Time spans are indicated in titles (0-1 h,1-2 h,2-3 h,3-4 h). Note that the stones in the middle affect a little the detected position of the fish, resulting in a more noisy location pattern of the green fish as compared to the others. D: Overall domains. A spatial region belongs to a particular fish if it occupies it for the most amount of time across the whole video. Black regions are never occupied (out side of the water level).

We found that **xyTracker** performed very well in this challenging circumstances (see Fig. 7). The inferred tracks of the individuals showed emergence of territories for each individual fish, which we confirmed to be correct by visual inspection of the video. In Fig. 7C the distribution of locations is calculated for sections of 1 hour for the first 4 hours. Note that over time a clear segregation of the 3 fish in space is seen. Overall, three clear domains emerged (Fig. 7D). Interestingly, fish 1 which was noticeable bigger, occupied the middle and largest region, relatively unchallenged, whereas fish 2 and fish 3 (of similar smaller size), initially contested fiercely for territories. This could be seen from the frequency of interaction (see Fig. 7 B) and was confirmed by visually inspecting the video. Taken together, **xyTracker** successfully tracked the individual fish under this challenging conditions and, importantly, did not confuse the identity of the fish as conventional tracking methods would be prone to. Note it would not have been possible to use **idTracker** successfully for this video because it would not cope with the challenging conditions described above.

### 3.4 Light avoidance task

We developed **xyTracker** specifically for reliable real-time tracking of group behavior and simultaneous visual stimulus presentation. For that we custom built an experimental setup where a LCD stimulus screen was placed below a transparent fish tank, while tracking was done using a camera mounted above the tank (Fig. 8A for an illustration). Since the camera observed fish from above, we tracked using infrared lightening and attached a visible light cutoff-filter to the camera so that the tracking was not distracted by the visual stimuli presentation (see Methods for a detailed description of the setup).

**Figure 8:**
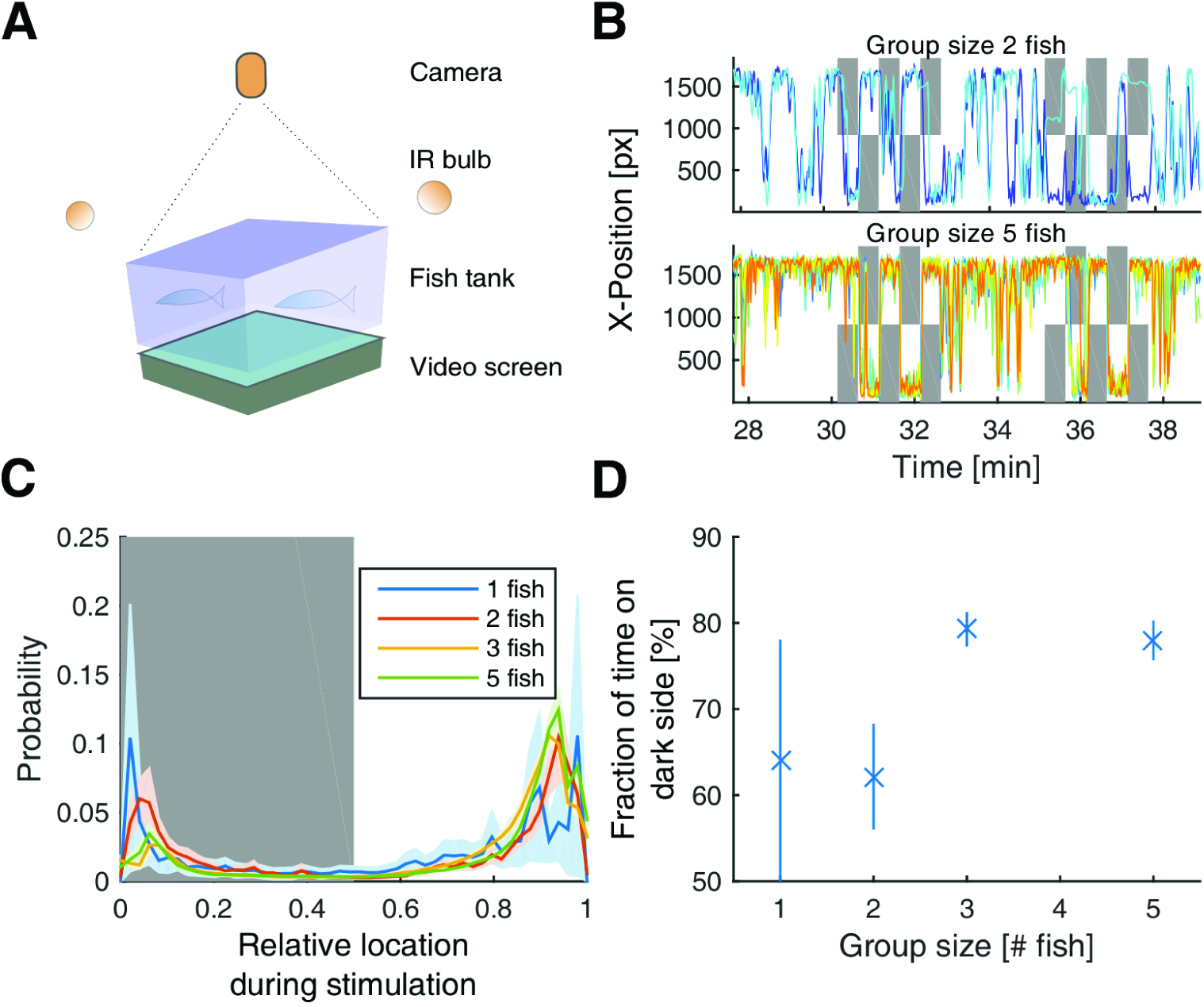
Group decision experiment: Avoidance of visual stimulus given from below. A: Experimental setup. The camera is installed above and tracks the fish illuminated by infrared bulbs. A video screen is install below the transparent tank. A cutoff filter is used to cut-off the visible light, so that the tracking is not hindered by the visual stimulus presentation. B: X-position over time of two groups of fish. Dark regions indicate stimulus presentation times. Stimulus is a half full-field white background that switches between left and right sides. C: Histogram of locations of fish, depending on the group size during stimulus presentation times. Note that fish prefer dark background over white backgrounds (white stimulus is indicated by gray). D: Choices of dark areas versus white background depend on the group size. The more individuals the higher is the fraction of time spend on the dark side.

To test whether visual stimuli presented on the screen from below affected fish behavior, we performed a simple light avoidance task. We asked how fish would react, when one half of the video screen showed an uniform white background, and the other half remained black. After selecting naive fish, we let them first adapt to the environment with visual stimulus turned off (at least^66^) for 30 min. After 30 min blank screen, we alternately displayed a white background on one half of the screen (either left or right) and reversed the position (either right or left) every 30 seconds. The stimulus changed 5 times and was then followed by a 150 seconds blank screen. This block repeated 5 times. To compare the behavior for different group sizes, we recorded and tracked individual fish for group sizes of 1 fish (3 experiments), 2 fish (3 experiments), 3 fish (5 experiments), and 5 fish (3 experiments), all with naive zebra fish.

The results are presented in Fig. 8. We found that zebra fish exhibit a strong avoidance behavior, usually leaving the region with light background quickly, similar to previously reported studies (done in fixed colored tanks) [38, 13]. Fish tend to rapidly swim into the dark region of the tank (see Fig. 8 B for example traces). This behavior was similar to the recently reported learning paradigm where fish had to learn the exit of a maze which was not illuminated from below [6]. Accordingly, during the stimulation the probability of localization was strongly biased towards the dark side Fig. 8 C. Interestingly, this light avoidance effect increased with the size of the group, so that larger groups (3 or 5 fish) showed stronger preference than small groups (Fig. 8 D). This group size effect, however, might be mainly due to the increased frequency of “freezing” when zebra fish are in groups of two or alone.

### 3.5 Closed-loop experiment

**xyTracker** was specifically designed to perform closed-loop experiments in behaving groups, where the presented stimulus depends on the collective behavior or on the particular location of individual members of the group.

To exemplify this type of experiments, we used the same setup as described in Fig. 8 A, but now let the stimulus depend on the actual location of individual fish. In particular, when the stimulation time was triggered of individual fish, we projected an image of a fish onto the screen at a position nearby the individuals. This “fish projections”, which we extracted using **xyTracker**, followed the position of a fish for a short amount of time. Moreover, based on the position, where a fish was located during the trigger event of the stimulus, the size of the projection was chosen: in one region, not stimulus was given (size 0), the second region, the projected image was of similar size as the fish, and the third regions, projected images tended to the twice as large as the fish (see example in Fig. 9 A).

**Figure 9:**
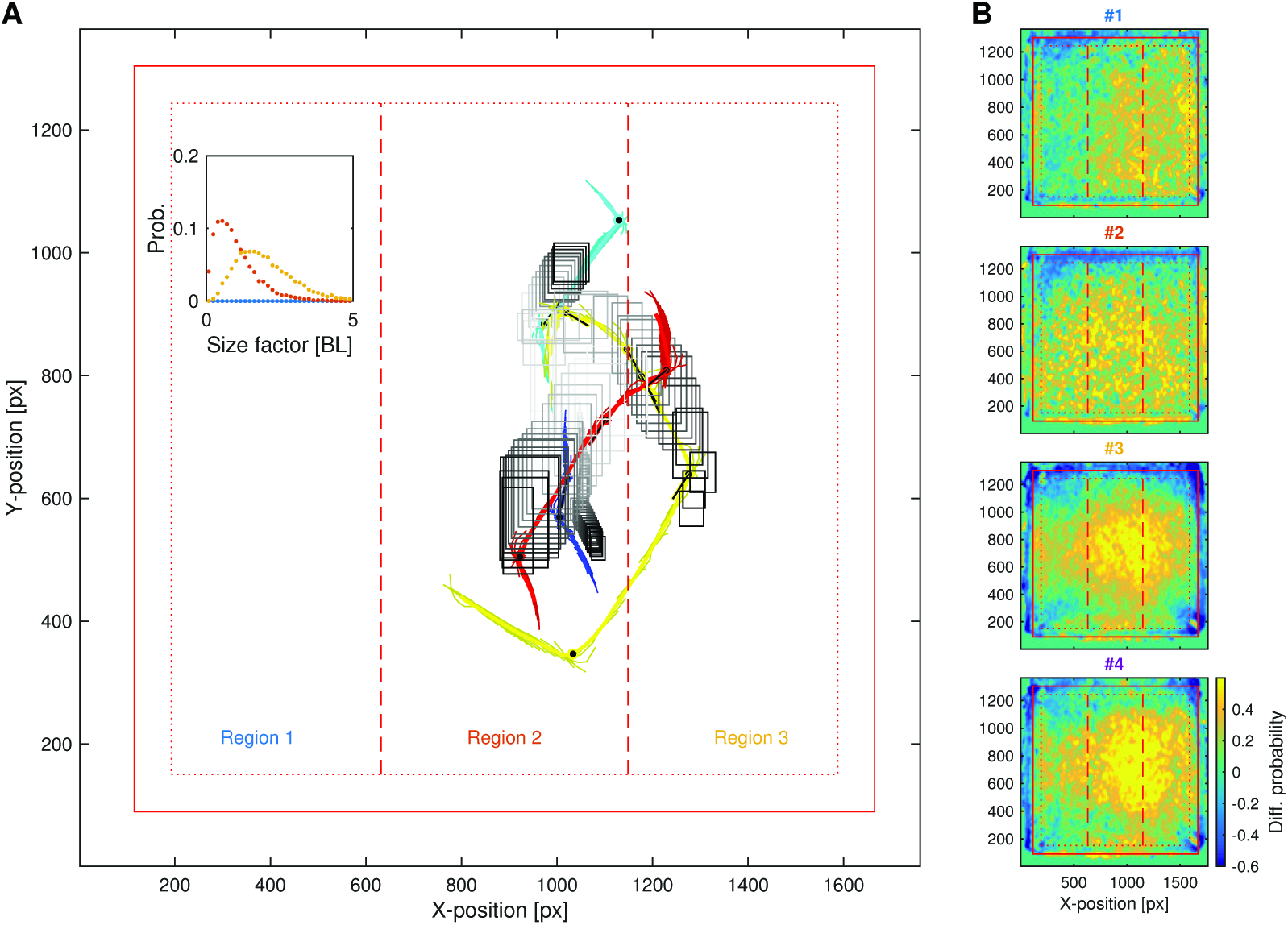
Closed-loop experiment. A: Stimulus presentation depends on the actual position of the tracked fish in real-time and last for about 500 ms. Solid lines indicate trajectories of fish for the past seconds. Rectangles indicate the bounding box of the fish projections which are generated based on the shapes of the fish during tracking. The gray scale of the bounding box outline indicates time: fish projections thus “follow” the fish. In the 3 regions, projections randomly occur with different size distributions. In region 1 and the borders (dotted red line) projections to not occur, so that fish can swim into these regions to avoid the projections. B: Difference probability maps for individual fish. For each position the fraction of time the velocity was larger than the median velocity minus the fraction of time it was lower was computed. Note that the velocity was generally higher in region 2 and 3 where the stimulus projections occurred. Moreover, fish learned to “relax” at the border region, where the probability of stimulation was zero. Thus the behavior of fish was clearly affected by the individual visual stimulation.

In detail, the position of individual fish was used and the extracted blob image (such as shown in e.g. in Fig. 3B) was projected onto the screen at randomly shifted positions relative to the actual fish position (on average 45 px offset; gamma distributed with Coefficient of Variation (CV) set to 0.5; orientation of the shift relative to the body center was uniformly distributed). The projections started at random times (on average 2 seconds, gamma distributed with CV set to 0.1) and lasted on average 500 ms (interval gamma distributed (CV) 0.1). Projections for individual fish had different colors. The size of the projections were scaled for each stimulus presentation time, where the size distribution depended on the region where the fish was at the time when the individual stimulus projection started (see inset in Fig. 9 A): In region 1 and the border regions (red dotted line), the size was set to 0, that is, no projections were shown. In region 2 average sizes were smaller than in region 3 (one average 1 body lengths (pixels) and 2 body lengths, respectively; gamma distributed with CV set to 0.5). At the border of the tank (5% of the length or width), no stimulus was presented, furthermore, size were set to 0 if the velocity was lower than a threshold.

We found that fish projections induced behavioral changes in individual fish, in particular, fish tried to escape from their following virtual projections (see example in Fig. 9 A). As a result of this escape, the fish velocity distribution was biased to regions 2 and 3, where the projections occurred randomly, and were slower in region 1 and in the border regions, where no projections occurred (see Fig. 9 B).

We thus conclude that it is possible to track and stimulate individual fish in a closed-loop fashion using **xyTracker**.

## 4 Discussion

We presented a new software platform for tracking of groups of behaving animals. Our system has a number of advancements and extensions over existing methods targeting the same application scope, in particular **idTracker** [45]. First, tracking occurs in an on-line manner, needing only a single pass through the video file or enabling tracking in real-time for closed-loop experiment. Second, we used a performance optimized implementation to achieve a > 30× speed-up using small amounts of memory, so that long and high-resolution videos with sizes of many gigabytes can still be tracked in an efficient manner. Third, our implementation is object-oriented and provides a convenient and extensible Matlab-interface. Forth, most parameters are self-tuned to the problem at hand, so that tracking will work out-of-the-box for most applications. In particular, the identity features used for recognizing the individuals are learned in an unsupervised and on-line manner (without need to specify features by hand), and are based simply on rotated image patches of the animals. Thus, these features contain all locally available image content, and therefore can be used and are informative for the tracking of any animal or moving object. In case of fish (or other elongated animals), our method further provides an more accurate estimate of the position than other comparable methods, because our approach includes a simple shape model, namely the mid line of the body. Thus, the position information of tracked animals is not only based on the center of a bounding box (which could even lie outside of the body, when the body bends during turns), but the exact position of specified part of the body, e.g. the head.

Many current multiple object tracking systems are developed specifically for the application to track humans [59, 58, 33]. The application of tracking humans in cluttered environment as specific challenges. Our application, the tracking of animals in a 2D arena, is more constrained. For instance, we assume that the environment, the camera, and the number of animals are fixed and (mostly) visible during the time of the experiment. One particular challenge in our situation is that, the appearance of individuals in an social group of animals, such as fish, are often highly similar. Moreover, when observed from above, their detected shape changes dramatically, because of body movement, bending and, direction of movement changes (rotations). Thus, a simple classifier for distinguishing the identity of individuals is easily distracted by the rotation and body deformations. The approach of [45] is to compute the histogram of two pixels’ intensity with given distance between the two pixels in the detected shape. While this feature is rotation invariant and informative about the identity, it is rather costly to compute (proportional to n2 where n is the number of pixels in the detection), does not account for deformations, and might neglect some useful information about the appearance available in the local image patch. We here solve the problem of rotation invariance in a very straightforward way: exploiting the elongated shape of animals and their asymmetry in the direction of movement (head front, tail back), we rotate the image patches to a reference orientation. The advantage of this direct rotation method is that all visual information for identity classification is still available. We found that even simple classification models could distinguish fish successfully using these rotated identity features.

The only limitation of the explicit rotation approach is the accuracy of the estimation of reference orientation. This reference rotation axis is inferred from the shape of the animals, thus in case of roughly circular objects, or animals with complicated symmetry, this reference axis might be difficult to estimate. However, we found that rotational invariance, and thus the explicit rotation, is not necessary for more powerful classifiers, such as CNN, which could be integrated into our object-based software platform in future^67^. Alternatively, our system could be extended to estimate the references axis from the movement direction instead of the geometric shape of the animals. This is not easily possible in the current implementation of **xyTracker**, because currently the rotation of the image patches is done independently from the tracking for increasing the speed and the modularity of the software code.

In computer vision approaches to tracking humans in cluttered environments, some methods also acquire an appearance model to improve the tracking results. A general framework to combine the tracking-by-detection with an appearance model is to use the multiple hypothesis tracking (MHT) approach [47, 30]. At each time step, MHT considers all possible association of detected objects with current tracks and builds a tree of all tracks. The method then performs an optimization to find the most likely tracks in the tree and thus can also distinguish and disentangle crossing events. We here developed a greedy version of MHT, based on a fast online computation of the shortest path through the track-tree. While very efficient, our approach does a shortest-path calculation without the constraint that detected objects belong only to one unique individual. If this constraint would be taken into account, the fast online computation would not be possible anymore, as it would need to be solved with a linear programming approach [3, 46]. However, for a smaller number of animals, the computational effort would probably still tractable, and thus integrating this constraint into **xyTracker** would be a worthwhile extension for the future.

Currently, our MHT approach might result in briefly overlapping trajectories for different animals. These overlaps tend to be short and negligible if enough appearance information is available. However, when the appearance information is more limited, e.g. when the available pixels per animals is small or lightening is challenging, so that individual animals become indistinguishable, our MHT method might result in a considerable number of overlapped tracks. We found that in this case, our explicit switching method outperforms the MHT approach. Under good conditions, however, the MHT approach usually performs better, mainly because all tracks are saved so that the switching of the track does not need to be inferred from the history, as in our explicit switching approach. In any case, since our software simultaneously calculates both approaches, resulting tracks can be compared and the one or the other can be used for further analysis. Moreover, **xyTracker** provides class probabilities for each frame that can be used to access the confidence about the current track identities.

Since our method switches track assignments in retrospect, assignments, in particular shortly after crossing events, might be momentarily wrong. While this post-hoc change is a drawback for online experiments, in practice, a momentarily wrong assignment is usually limited to a short period after crossings where animals are in close proximity anyway. Also, a track switching in retrospect might be unavoidable and acceptable if it is then ensured that the identity of the tracks are corrected in the following. We showed exemplary how to build a closed-loop experiment to perturb the behavior of individual fish by locally targeted presentation of small visual stimuli. Thus, in our setup, one could replicate experiments where a robotic fish was employed [7], using instead projected images of fish. **xyTracker** is based on the PsychToolbox for stimulus generation and thus easily integrates with OpenGL features for a wide range of stimulus capabilities. In principle, also any other stimulus generation system could be integrated, e.g. for generating more advanced virtual reality environments, which than could be used to test models of collective response to perturbations in experiments [16].

In conclusion, for tracking animals for behavioral studies of social groups it is essential to ensure the identity of individual animals for a long period of time. We showed that our system can stably follow individual fish for hours. Furthermore, we showed how **xyTracke**r's online tracking capabilities can be used to perform challenging closed-loop group learning experiments.

## Footnotes

1 **xyTracker** is available for download at https://www.github.com/maljoras/xyTracker.

2 That is, almost all parameters will be estimated automatically and thus the system should work out-of-the-box in most cases.

3 We found that for our videos a simple uniform gray-scale mapping of RGB values is enough (that is taking the RGB color mean). However, **xyTracker** supports an object enhancing mapping of RGB values into gray scale pictures based on a linear fisher discriminant projection (Option **useScaledFormat**). This could be useful for similar colored fish that swim in front of a diffierent colored background, which intensity level is, however, similar to that of the animals.

4 The **xyTracker** also supports other method such as k-nearest neighbor background model from the OpenCV toolbox (option `useKNN`). Although accuracy increased, it turned out to be a bottleneck in computational speed.

5 Controlled by setting the option **`detector.inverted`** in **xyTracker**.

6 Option **`detector.nskip`**.

7 Option `detector.history`

8 Otherwise immobile animals will become inseparable from the background with time.

9 Option `detector.adjustThresScale`. Noisiness of the detections can be checked when displaying the mask during the tracking (by setting the option `display.BWImg`)

10 Options `bodylength` and `bodywidth`

11 However, when calculating the center line distance below, we take the minimal distance between center-line points in straight or reversed order, see Table 1.

12 This is particular useful for tracking fish. However, one could use other references for other animals if needed. For instance, the orientation of the major axis of the fitted ellipse for oval shaped animals.

13 Option `tracks.kalmanFilterPredcition`

14 We set in the results γ = (10; 2; 5; 1; 0; 0; 0), compare to Table 1. Options in **xyTracker** are of the form cost.feature, e.g. `cost.Location` or `cost.Overlap`

15 Note that this cost-weightening could be done using more sophisticated updates, e.g. Bayesian methods or Kalman filters. However, we here choose a simple running mean for the sake of speed.

16 Option `tracks.costOfNonAssignment`

17 Note that we currently assume that animals never remain invisible for a long time. If this was the case, **xyTracker** will produce noise detections for the lost tracks and tracking results generally will deteriorate. Only in a future version, **xyTracker** might support the deletion an creation of tracks.

18 We define “far” relative to the length of the body *L*.

19 We assume that the frame rate of the video is high enough so that bounding boxes typically overlap between frames

20 Option `useDagResults` sets DAG to default

21 The size of the part from the head to tail can be adjusted if necessary and is only intended to cut off the movement in the tails of fish and might not be needed when tracking other animals

22 E.g. a convolutional net by implementing or overloading the `FishBatchClassifier` class

23 It derives from the class `xy.core.BatchClassifier`.

24 Option `classifier.timeToInit`

25 However, even if some crossings occur, initialization will start after several hundred frames anyway. Thus **xyTracker** works best when animals do no cluster together and move freely in the beginning.

26 Option `classifier.npca`

27 Option `classifier.nlfd`

28 Property `reasonableThres` in the `xy.core.ClassProbHistory` class.

29 Property taulambda in the `xy.core.ClassProbHistory class`.

30 This parameter is currently set by hand and not estimated based on the application context. In our videos, a value of 70 frames worked well. This parameter can be set by adjusting the `classifier.timeForUniqueUpdate` option.

31 Option `classifier.timeForSingleUpdate`

32 Option `classifier.tau`

33 *ρ*_forced_, `option classifier.forcedUpdateProbThres`. In detail, we re-assigned the tracks according to the results of the classifier prediction, when the maximal class probability of the permuted tracks is smaller than *ρ*_forced_ *p*_max_, where *p*_max_ is a running average (with time constant `tracks.costtau`) of the highest class probability in each frame, while the mean of the class probabilities has to be higher than *_ρ_*_identity_ *p*_max_ (*ρ*_identity_ corresponds to option **tracks.probThresForIdentity**). Alternatively, to allow for accurate learning, it is also updated if *p*mean - *p*other < *p*diffother, where *p*_mean_ is the running average of the mean probability per frame and *p*_other_ is the running average of the maximal “other” class probability, that is, if the tracks would be assigned to any other identity and *p*_diffother_ a parameter (option `classifier.minOtherProbDiff`). In this way it is ensured that meaningful identities have been learned before using the appearance model to reassign the identity of the tracks.

34 option `classifier.allSwitchProbThres`. In detail, if the average (positive) difference probability of the predicted identities minus the original identities is larger than *ρ*_allswitch_ *p*_max_ (*ρ*_allswitch_ corresponds to option `classifier.allSwitchProbThres`), then the reassignment of tracks is considered in the same way as is handled after crossings (see below).

35 Option `tracks.crossBoxLengthScale`

36 The position of a invisible track is the position when it was last visible.

37 Option `classifier.crossCostThresScale`

38 Option `classifier.timeAfterCrossing`

39 Option `classifier.reassignProbThres`

40 Option `classifier.timeAfterCrossing`

41 Option `dag.probScale`

42 In fact we optional (option `tracks.centerLinePrediction`) shift the location (usually the center of the center line points) along the center line points according to the current track velocity (which is estimated in a leaky average fashion (option `classifier.clpMovAvgTau`)). One could also include a predicted location, e.g. using Kalman filter (option `tracks.kalmanFilterPredcition`), but we found that this yields inferior results if the frame rate is reasonable high.

43 This can be done without additional noticeable computational cost because of the mutli-threading structure of the code.

44 Set default method by option `tracks.useDagResults`. See also methods `getDagTrackingResults` or `getSwitchBasedTrackingResults`

45 We use the DAG method in some case to compute the shortest path only for the recent history to disentangle the identities in retrospect for the SWB method.

46 DAG-based identity prediction for the stimulus presenter can be chosen by setting the option `stimulus.usePredIdentityId`

47 Base class `xy.stimulus.Presenter`. See e.g. `xy.stimulus.PresenterFlash` for an example how to overload the base class.

48 **xyTracker** is available for download at https://www.github.com/maljoras/xyTracker.

49 http://opencv.org/

50 Version 3: set option `useMex` to 1. Version 2: set option `useMex` to 0 and `useOpenCV` 1. Version 3: set both options to 0.

51 https://github.com/kyamagu/mexopencv

52 For instance, only version 3 currently supports the “center line” computations.

53 This can be tested by calling `fish.Tracker.runTest`(100)

54 One possible workaround of the memory problem would be to divide the video into many small parts, track the parts, and recognize the same individuals across parts. However, we have not tried this, since given the > 30× speed up advantage of **xyTracker**, the tracking would need at least 9 days on this video with `idTracker`.

55 for now only version (3) supports grabbing, although adding this feature to the other versions would be straightforward, when using a OpenCV / Matlab vision toolbox compatible camera

56 https://www.ptgrey.com/

57 https://ffmpeg.org/

58 The exact time point when the frame was grabbed is nevertheless known and used by the tracking system. The frame rate of the background video recording is independent from the tracking frame rate.

59 see https://www.github.com/maljoras/xyTracker

60 The run test will generate Fig. 6.

61 It is recommended to start Matlab with the switch `-nodesktop`

62 Or set later with `T.setOpts(opts)`, or directly accessing the `T.opts` property. Note, however, that some options (mostly those with no additional “.” in their name) can only be given at the beginning during instantiation of the **xyTracker** object, since they are needed for initialization. In particular, the number of tracked animals, i.e. `nindiv`, should be given if known, otherwise it will be estimated automatically.

63 The structure has to be the last element with fields corresponding to the name of the options.

64 On our videos, the **idTracker** method performed very poorly, because the lightening condition were more challenging.

65 Size 45 cm×45 cm, water depth about 10 cm

66 most groups were put into the experimental setup the day before the experiment started and thus adapted at least 12h.

67 However, to learn these more powerful classifiers more data will be needed.

## References

1. M. B. Ahrens, J. M. Li, M. B. Orger, D. N. Robson, A. F. Schier, F. Engert, and R. Portugues. Brain-wide neuronal dynamics during motor adaptation in zebrafish. Nature, 485(7399):471–477, 2012.

2. L. Al-Imari and R. Gerlai. Sight of conspecifics as reward in associative learning in zebrafish (danio rerio). Behavioural brain research, 189(1):216–219, 2008.

3. C. Amit Kumar K, D. Delannay, and C. De Vleeschouwer. Iterative hypothesis testing for multi-object tracking in presence of features with variable reliability. arXiv preprint arXiv:1509.00313, 2015.

4. A. Amsterdam and N. Hopkins. Mutagenesis strategies in zebrafish for identifying genes involved in development and disease. Trends in Genetics, 22(9):473–478, 2006.

5. D. J. Anderson and P. Perona. Toward a science of computational ethology. Neuron, 84(1):18–31, 2014.

6. R. Aoki, T. Tsuboi, and H. Okamoto. Y-maze avoidance: An automated and rapid associative learning paradigm in zebrafish. Neuroscience research, 91:69–72, 2015.

7. M. Aureli, F. Fiorilli, and M. Porfiri. Portraits of self-organization in fish schools interacting with robots. Physica D: Nonlinear Phenomena, 241(9):908–920, 2012.

8. M. Ballerini, N. Cabibbo, R. Candelier, A. Cavagna, E. Cisbani, I. Giardina, V. Lecomte, A. Orlandi, G. Parisi, A. Procaccini, et al. Interaction ruling animal collective behavior depends on topological rather than metric distance: Evidence from a field study. Proceedings of the national academy of sciences, 105(4):1232–1237, 2008.

9. J. Bergstra, O. Breuleux, F. Bastien, P. Lamblin, R. Pascanu, G. Desjardins, J. Turian, D. Warde-Farley, and Y. Bengio. Theano: a cpu and gpu math expression compiler. In Proceedings of the Python for scientific computing conference (SciPy), volume 4, page 3. Austin, TX, 2010.

10. I. H. Bianco and F. Engert. Visuomotor transformations underlying hunting behavior in zebrafish. Current Biology, 25(7):831–846, 2015.

11. C. M. Bishop. Pattern recognition. Machine Learning, 2006.

12. S. S. Blackman. Multiple hypothesis tracking for multiple target tracking. *Aerospace and Electronic Systems Magazine*, IEEE, 19(1):5–18, 2004.

13. R. E. Blaser and D. B. Rosemberg. Measures of anxiety in zebrafish (danio rerio): dissociation of black/white preference and novel tank test. PLoS One, 7(5):e36931, 2012.

14. B. Brembs and M. Heisenberg. The operant and the classical in conditioned orientation of drosophila melanogaster at the flight simulator. Learning & Memory, 7(2):104–115, 2000.

15. D. S. Calovi, U. Lopez, S. Ngo, C. Sire, H. Chaté, and G. Theraulaz. Swarming, schooling, milling: phase diagram of a data-driven fish school model. New Journal of Physics, 16(1):015026, 2014.

16. D. S. Calovi, U. Lopez, P. Schuhmacher, H. Chaté, C. Sire, and G. Theraulaz. Collective response to perturbations in a data-driven fish school model. Journal of The Royal Society Interface, 12(104):20141362, 2015.

17. N. Chenouard, I. Smal, F. De Chaumont, M. Maška, I. F. Sbalzarini, Y. Gong, J. Cardinale, C. Carthel, S. Coraluppi, M. Winter, et al. Objective comparison of particle tracking methods. Nature methods, 11(3):281, 2014.

18. T. H. Cormen. Introduction to algorithms. MIT press, 2009.

19. I. D. Couzin, J. Krause, N. R. Franks, and S. A. Levin. Effective leadership and decisionmaking in animal groups on the move. Nature, 433(7025):513–516, 2005.

20. L. J. Cox and S. L. Hingorani. An efficient implementation of reid's multiple hypothesis tracking algorithm and its evaluation for the purpose of visual tracking. *Pattern Analysis and Machine Intelligence*, IEEE Transactions on, 18(2):138–150, 1996.

21. A. I. Dell, J. A. Bender, K. Branson, I. D. Couzin, G. G. de Polavieja, L. P. Noldus, A. Pérez-Escudero, P. Perona, A. D. Straw, M. Wikelski, et al. Automated image-based tracking and its application in ecology. Trends in Ecology & Evolution, 29(7):417–428, 2014.

22. M. B. Dillencourt, H. Samet, and M. Tamminen. A general approach to connected-component labeling for arbitrary image representations. Journal of the ACM (JACM), 39(2):253–280, 1992.

23. J. Duchi, E. Hazan, and Y. Singer. Adaptive subgradient methods for online learning and stochastic optimization. The Journal of Machine Learning Research, 12:2121–2159, 2011.

24. R. E. Engeszer, L. B. Patterson, A. A. Rao, and D. M. Parichy. Zebrafish in the wild: a review of natural history and new notes from the field. Zebrafish, 4(1):21–40, 2007.

25. T. Fukunaga, S. Kubota, S. Oda, and W. Iwasaki. Grouptracker: Video tracking system for multiple animals under severe occlusion. Computational biology and chemistry, 57:39–45, 2015.

26. J. Gautrais, F. Ginelli, R. Fournier, S. Blanco, M. Soria, H. Chaté, and G. Theraulaz. Deciphering interactions in moving animal groups. PLoS Comput Biol, 8(9):e1002678, 2012.

27. R. Gerlai. A small fish with a big future: zebrafish in behavioral neuroscience. Reviews in the Neurosciences, 22(1):3–4, 2011.

28. G. E. Hinton, N. Srivastava, A. Krizhevsky, I. Sutskever, and R. R. Salakhutdinov. Improving neural networks by preventing co-adaptation of feature detectors. arXiv preprint arXiv:1207. 0580, 2012.

29. M. Kabra, A. A. Robie, M. Rivera-Alba, S. Branson, and K. Branson. Jaaba: interactive machine learning for automatic annotation of animal behavior. nature methods, 10(1):64–67, 2013.

30. C. Kim, F. Li, A. Ciptadi, and J. M. Rehg. Multiple hypothesis tracking revisited. In Proceedings of the IEEE International Conference on Computer Vision, pages 4696–4704, 2015.

31. M. Kleiner, D. Brainard, D. Pelli, A. Ingling, R. Murray, C. Broussard, et al. Whats new in psychtoolbox-3. Perception, 36(14):1, 2007.

32. H. W. Kuhn. The hungarian method for the assignment problem. Naval research logistics quarterly, 2(1-2):83–97, 1955.

33. L. Leal-Taixé, A. Milan, I. Reid, S. Roth, and K. Schindler. Motchallenge 2015: Towards a benchmark for multi-target tracking. arXiv preprint arXiv:1504.01942, 2015.

34. Y. LeCun, L. Bottou, Y. Bengio, and P. Hanner. Gradient-based learning applied to document recognition. Proceedings of the IEEE, 86(11):2278–2324, 1998.

35. C. M. Lindeyer and S. M. Reader. Social learning of escape routes in zebrafish and the stability of behavioural traditions. Animal Behaviour, 79(4):827–834, 2010.

36. U. Lopez, J. Gautrais, I. D. Couzin, and G. Theraulaz. From behavioural analyses to models of collective motion in fish schools. Interface focus, page rsfs20120033, 2012.

37. R. Lukeman, Y.-X. Li, and L. Edelstein-Keshet. Inferring individual rules from collective behavior. Proceedings of the National Academy of Sciences, 107(28):12576–12580, 2010.

38. C. Maximino, T. M. de Brito, C. A. G. de Mattos Dias, A. Gouveia, and S. Morato. Scototaxis as anxiety-like behavior in fish. Nature protocols, 5(2):209–216, 2010.

39. G. McLachlan. Discriminant analysis and statistical pattern recognition, volume 544. John Wiley & Sons, 2004.

40. D. P. Mersch, A. Crespi, and L. Keller. Tracking individuals shows spatial fidelity is a key regulator of ant social organization. Science, 340(6136):1090–1093, 2013.

41. N. Miller and R. Gerlai. From schooling to shoaling: patterns of collective motion in zebrafish (danio rerio). PLoS One, 7(11):e48865, 2012.

42. K. P. Mueller and S. C. Neuhauss. Automated visual choice discrimination learning in zebrafish (danio rerio). Journal of integrative neuroscience, 11(01):73–85, 2012.

43. V. Nair and G. E. Hinton. Rectified linear units improve restricted boltzmann machines. In Proceedings of the 27th International Conference on Machine Learning (ICML-10), pages 807–814, 2010.

44. R. F. Oliveira. Mind the fish: zebrafish as a model in cognitive social neuroscience. Frontiers in neural circuits, 2014.

45. A. Péerez-Escudero, J. Vicente-Page, R. C. Hinz, S. Arganda, and G. G. de Polavieja. idtracker: tracking individuals in a group by automatic identification of unmarked animals. Nature methods, 11(7):743–748, 2014.

46. H. Pirsiavash, D. Ramanan, and C. C. Fowlkes. Globally-optimal greedy algorithms for tracking a variable number of objects. In Computer Vision and Pattern Recognition (CVPR), 2011 IEEE Conference on, pages 1201–1208. IEEE, 2011.

47. D. B. Reid. An algorithm for tracking multiple targets. Automatic Control, IEEE Transactions on, 24(6):843–854, 1979.

48. K. E. Severi, R. Portugues, J. C. Marques, D. M. OMalley, M. B. Orger, and F. Engert. Neural control and modulation of swimming speed in the larval zebrafish. Neuron, 83(3):692–707, 2014.

49. Y. Shemesh, Y. Sztainberg, O. Forkosh, T. Shlapobersky, A. Chen, and E. Schneidman. High-order social interactions in groups of mice. Elife, 2:e00759, 2013.

50. M. Sison and R. Gerlai. Associative learning in zebrafish (danio rerio) in the plus maze. Behavioural brain research, 207(1):99–104, 2010.

51. R. Spence. Zebrafish ecology and behaviour. Zebrafish Models in Neurobehavioral Research, pages 1–46, 2011.

52. R. Spence, G. Gerlach, C. Lawrence, and C. Smith. The behaviour and ecology of the zebrafish, danio rerio. Biological Reviews, 83(1):13–34, 2008.

53. R. Spence and C. Smith. Male territoriality mediates density and sex ratio effects on oviposition in the zebrafish, danio rerio. Animal Behaviour, 69(6):1317–1323, 2005.

54. M. D. Suboski, S. Bain, A. E. Carty, L. M. McQuoid, M. I. Seelen, and M. Seifert. Alarm reaction in acquisition and social transmission of simulated-predator recognition by zebra danio fish (brachydanio rerio). Journal of Comparative Psychology, 104(1):101, 1990.

55. S. V. Viscido, J. K. Parrish, and D. Grϋnbaum. Individual behavior and emergent properties of fish schools: a comparison of observation and theory: Emergent properties of complex marine systems: a macroecological perspective. Marine ecology. Progress series, 273:239–249, 2004.

56. B. Wang, G. Wang, K. L. Chan, and L. Wang. Tracklet association by online target-specific metric learning and coherent dynamics estimation. arXiv preprint arXiv:1511.06654, 2015.

57. A. Weissbrod, A. Shapiro, G. Vasserman, L. Edry, M. Dayan, A. Yitzhaky, L. Hertzberg, O. Feinerman, and T. Kimchi. Automated long-term tracking and social behavioural phenotyping of animal colonies within a semi-natural environment. Nature communications, 4, 2013.

58. H. Yang, L. Shao, F. Zheng, L. Wang, and Z. Song. Recent advances and trends in visual tracking: A review. Neurocomputing, 74(18):3823–3831, 2011.

59. A. Yilmaz, O. Javed, and M. Shah. Object tracking: A survey. Acm computing surveys (CSUR), 38(4):13, 2006.

